# Constructing synthetic-protein assemblies from *de novo* designed 3_10_ helices

**DOI:** 10.1101/2021.12.11.471898

**Authors:** Prasun Kumar, Neil G. Paterson, Jonathan Clayden, Derek N. Woolfson

## Abstract

Compared with the iconic α helix, 3_10_ helices occur much less frequently in protein structures. The different 3_10_-helical parameters lead to energetically less favourable internal energies, and a reduced tendency to pack into defined higher-order structures. Consequently, in natural proteins, 3_10_ helices rarely extend past 6 residues, and do not form regular supersecondary, tertiary, or quaternary interactions. Here, we show that despite their absence in nature, synthetic protein-like assemblies can be built from 3_10_ helices. We report the rational design, solution-phase characterisation, and an X-ray crystal structure for water-soluble bundles of 3_10_ helices with consolidated hydrophobic cores. The design uses 6-residue repeats informed by analysing natural 3_10_ helices, and incorporates aminoisobutyric acid residues. Design iterations reveal a tipping point between α-helical and 3_10_-helical folding, and identify features required for stabilising assemblies in this unexplored region of protein-structure space.

---

The α helix is one of the fundamental constants of biology: it is a cornerstone of structural biology^1^, a model for protein-folding studies^2,3^, a workhorse in protein design^4,5^, and a scaffold for displaying functional moieties in protein engineering and biotechnology^6,7^. Largely, this is because it is highly defined conformationally: it has a narrow range of backbone-torsion or Ramachandran angles^8^, which leads to tight helical parameters, **Fig 1**; and stabilising intrahelical backbone hydrogen bonding between residues ‘i’ and ‘i+4’ along the polypeptide chain, CO_i_→NH_i+4_. Put another way, energetically, the α helix sits in a narrow and deep free-energy well^9^. As a result, the α helix can be considered a discrete building block for protein structures and assemblies, which accounts for its pre-eminence and biology and exploitation in protein design and engineering.

**Fig. 1:**
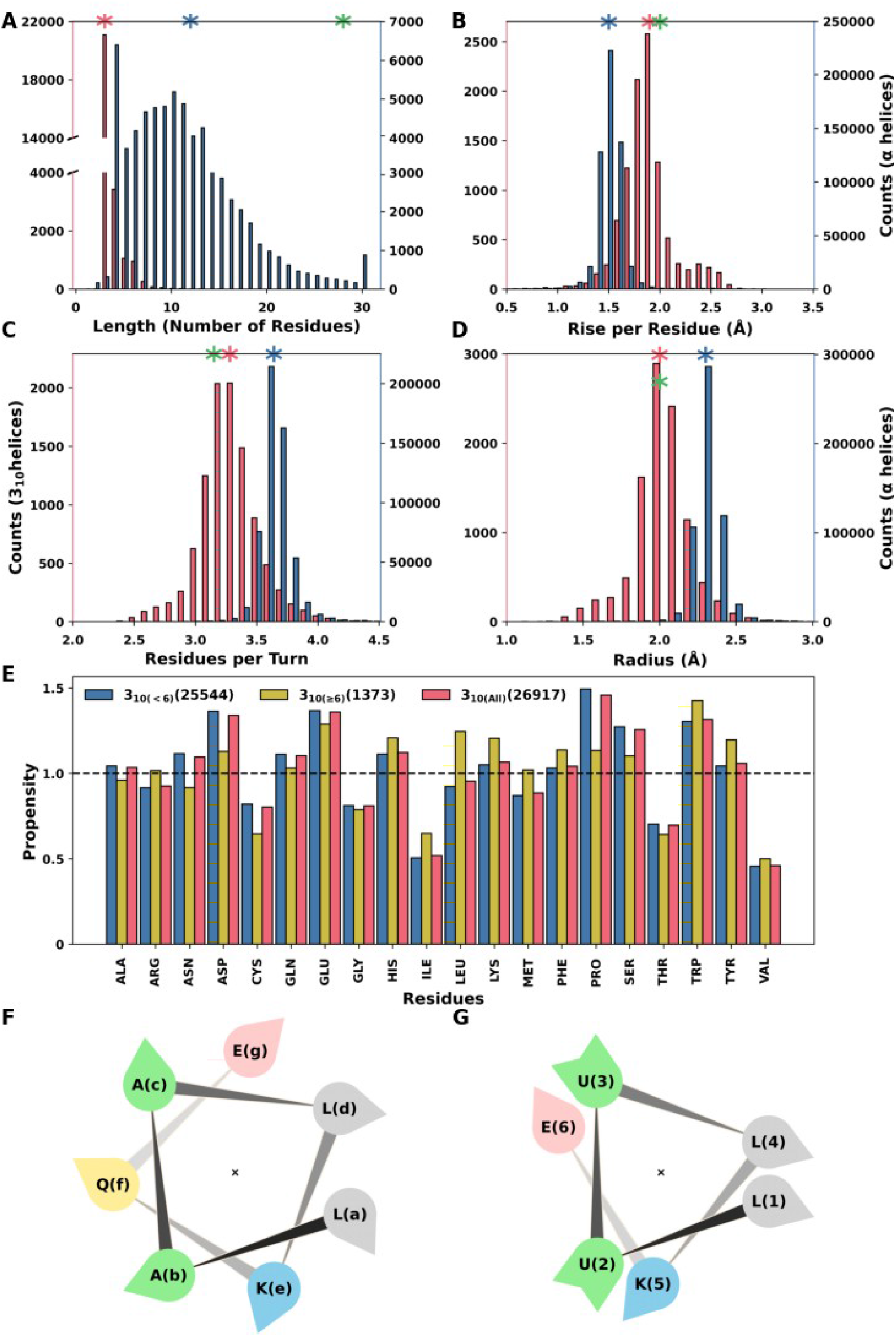
Analysis of the PDB and design principles for 3_10_ helices. Helical parameters for 3_10_ helices (red) and α helices (blue) calculated from the PDB: (**A**) helix length; (**B**) rise per residue; (**C**) residues per turn; and (**D**) radius. Mean values are marked with red and blue asterisks. The green asterisks show the mean values from the new quaternary structure of 3_10_ helices reported here. For panel **A**, all helices over 29 residues long are plotted as a single bar at 30 residues; the longest α helix found had 97 residues. (**E**) Propensity values of the 20 standard amino acids in 3_10_ helices, divided into two categories for helices shorter than 6 residues long (3_10 (<6)_; blue) or greater than 5 residues long (3_10 (≥6)_; orange). (**F** and **G**) Helical-wheel diagrams for a *de novo* 7-residue sequence repeat on an α helix (**F**), and a *de novo* 6-residue repeat on a 3_10_ helix (**G**).

Nonetheless, other helical peptide conformations do exist near to the α-helical region of Ramachandran space—namely, 3_10_-helices and π-helices^10^. These have different helical parameters leading to tighter and looser helical structures with CO_i_→NH_i+3_ and CO_i_→NH_i+5_ hydrogen-bonding patterns, respectively. They lie on the rim of the α-helical free-energy well and are, thus, thermodynamically less stable. Consequently, it is not surprising that these potentially accessible polypeptide conformations occur less commonly in natural proteins. Indeed, we surveyed a non-redundant set of protein structures from the RCSB Protein Dank (PDB)^11^ and found that while 33% of the >2 million residues were located within α helices, only 4% were within 3_10_ helices, and 0.5% within π helices, **Fig. 1** and **Table S7.1**. Moreover, the 3_10_ and π helices were much shorter than the α helices, **Fig. 1A** and **Fig. S1.1**. Furthermore, we found no examples in proteins where consolidated packing arrangements— i.e., supersecondary, tertiary and quaternary interactions—are formed exclusively by these alternative helices. By contrast, such structures based on α helices are widely catalogued^12-15^. Evidently, nature appears not to use 3_10_ or π helices to construct higher-order peptide and protein assemblies in aqueous media. This raises the question of whether such objects can nonetheless exist, and the challenge of whether chemists might be able to make them.

One group of natural peptides do commonly adopt 3_10_-helical conformations; namely, the membrane-active fungal metabolites known as the peptaibols^16,17^. These short peptides are rich in α,α-disubstituted amino acids, in particular α-aminoisobutyric acid (Aib, U)^18-24^, which favour tighter helical turns characteristic of 3_10_ helices^25-28^. However, the formation of 3_10_ helices appears to fall off with lengths >9 residues, where α-helix formation becomes favoured^29^. Moreover, while oligomers of Aib have been used as functional synthetic models of membrane-penetrating proteins such as G-protein coupled receptors^30^ and rhodopsin^31^, peptide solubility in water is lowered by increased content of hydrophobic Aib. Polar and charged groups can be added to increase water solubility of 3_10_-helical peptides, but atomistic structures remain elusive^32,33^.

Thus, whilst natural peptide and protein chains do access 3_10_-helical conformations, these tend not to propagate beyond short stretches, and they do not facilitate higher-order tertiary and quaternary interactions. Furthermore, the folding and assembly of 3_10_ helices appears to be even more limited in aqueous solution. We hypothesised that if we could design a synthetic (*de novo*) sequence to form an amphipathic 3_10_ helix of sufficient length—*i.e*., one with distinct hydrophobic and polar faces—it could be stabilised by helix-helix interactions to form a 3_10_-helical bundle. Here, we present the design and high-resolution structure of such a bundle.

## Rational design and characterisation of α- and 3_10_-helical bundles

We took a rational design approach to generate repeat sequences aimed at forming long 3_10_ helices in aqueous solution. Our starting point was the aforementioned set of protein structures harvested from the PDB. Although 3_10_ helices are fewer in number and shorter in length than α helices, **Fig. 1A**, the data set contained sufficient examples to calculate propensities for each of the 20 proteinogenic residues to adopt this conformation compared with any other state, **Fig. 1B** and **Table S7.1**. These data indicated that for 3_10_ helices of ≥6 residues, glutamate (Glu, E), lysine (Lys, K), leucine (Leu, L) and tryptophan (Trp, W) had preferences for 3_10_-helical secondary structures.

It is well understood that amphipathic α-helical peptides can be stabilised by association into coiled-coil bundles, and design principles for these are well established^4,5,34^. Therefore, we began our design process with a control α-helical bundle using the above amino-acid palette plus alanine (Ala, A). Amphipathic α helices can be encoded by placing hydrophobic residues 3 and 4 residues apart, as the average 3.5-residue spacing closely matches the 3.6 residues per turn of the helix, **Fig. 1F**. Based on past experience^35^, we designed the 7-residue repeat, E-L-A-A-L-K-X, where X can be any amino acid. We repeated this 4 times to make peptide PK-1 (systematically named CC-TypeN-L_a_L_d_), **Table 1**.

**Table 1:**
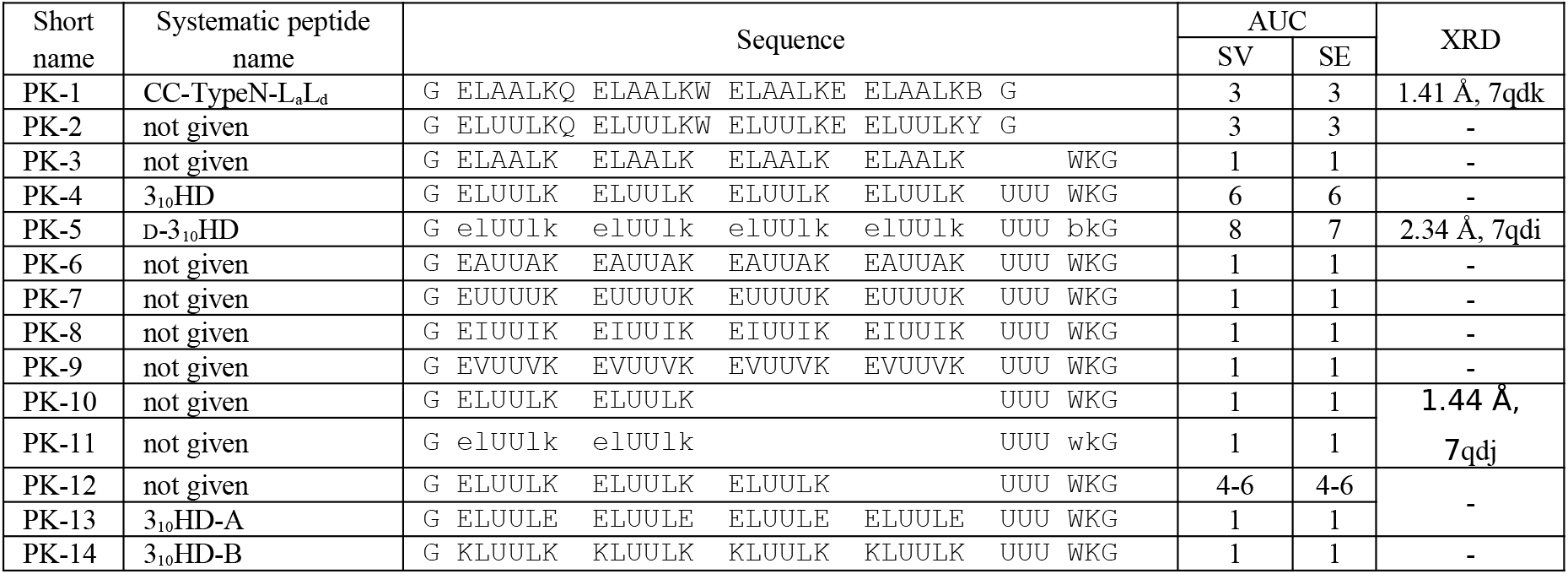
De novo designed peptide sequences. Standard amino acids are represented by their one letter codes, and U denotes Aib and B 4-bromo-l-phenylalanine. D-Amino acids are in lowercase. For the analytical ultracentrifugation (AUC) sedimentation velocity (SV) and sedimentation equilibrium (SE), the numbers indicate the oligomeric states determined from these experiments. Resolution (Å) and PDB identifiers are given for structures solved by X-ray diffraction (XRD). Peptides were made by solid-phase peptide synthesis and confirmed by mass spectrometry, Supplementary Information section 2.

Circular dichroism (CD) spectroscopy showed that PK-1 adopted a thermally stable α helix in solution, **Fig. 2A** and **Fig S3.1**, and analytical ultracentrifugation (AUC) indicated that it formed a single trimeric species, **Fig. 2A** and **Fig S4.1**. We crystallised and determined an X-ray structure for PK-1, which revealed a parallel, trimeric, α-helical, coiled-coil bundle, **Fig. 3A** and **Tables S7.2&3**, consistent with foregoing coiled-coil design principles^34-36^.

**Fig. 2:**
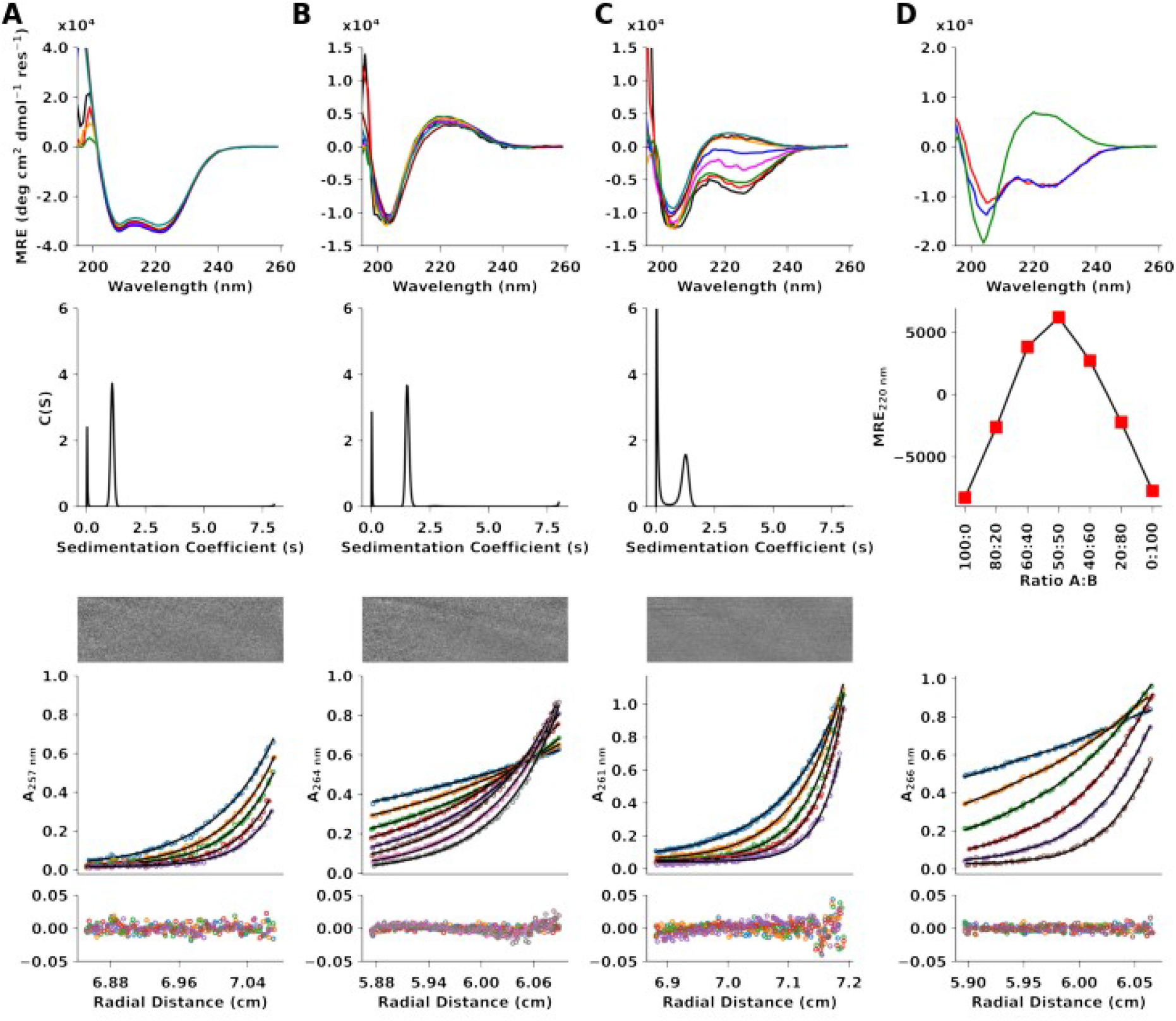
Biophysical characterization of *de novo* designed peptides. (**A – C**) Data for peptides PK-1 (**A**), PK-4 (3_10_HD) (**B**), and PK-12 (**C**), from top to bottom: CD spectra recorded at 5 °C; SV c(s) distributions; and SE plots. Conditions for CD experiments: PBS, pH 7.4, at peptide concentrations of 5 μM (black), 10 μM (red), 25 μM (green), 50 μM (magenta), 100 μM (blue), 200 μM (orange), 500 μM (maroon), 1000 μM (light green). AUC conditions: 20 °C, PBS, pH 7.4, 100 μM peptide concentration. Rotor speeds for SV: 60K RPM; Rotor speed for SE in panel **A** and **C**: 44k (blue), 48k (orange), 52k (green), 56k (red) and 60k (purple) RPM; Rotor speed for SE for panel **B**: 15k (blue), 18k (orange), 21k (green), 24k (red), 27k (purple), 30k (brown), 33k (pink) and 36k (grey) RPM. (**D**) Heteromeric 3_10_ design (3_10_HD-AB), from top to bottom: CD spectra at 5 °C, 100 μM peptide concentrations for the acidic (PK-13; red), basic (PK-14; blue) peptides and mixture (green); Job plot for the mixture of acidic and basic peptides; SE plots for the mixture. Rotor speed for SE for panel **D**: 18k (blue), 24k (orange), 30k (green), 36k (red), 42k (purple) and 48k (brown) RPM.

**Fig. 3:**
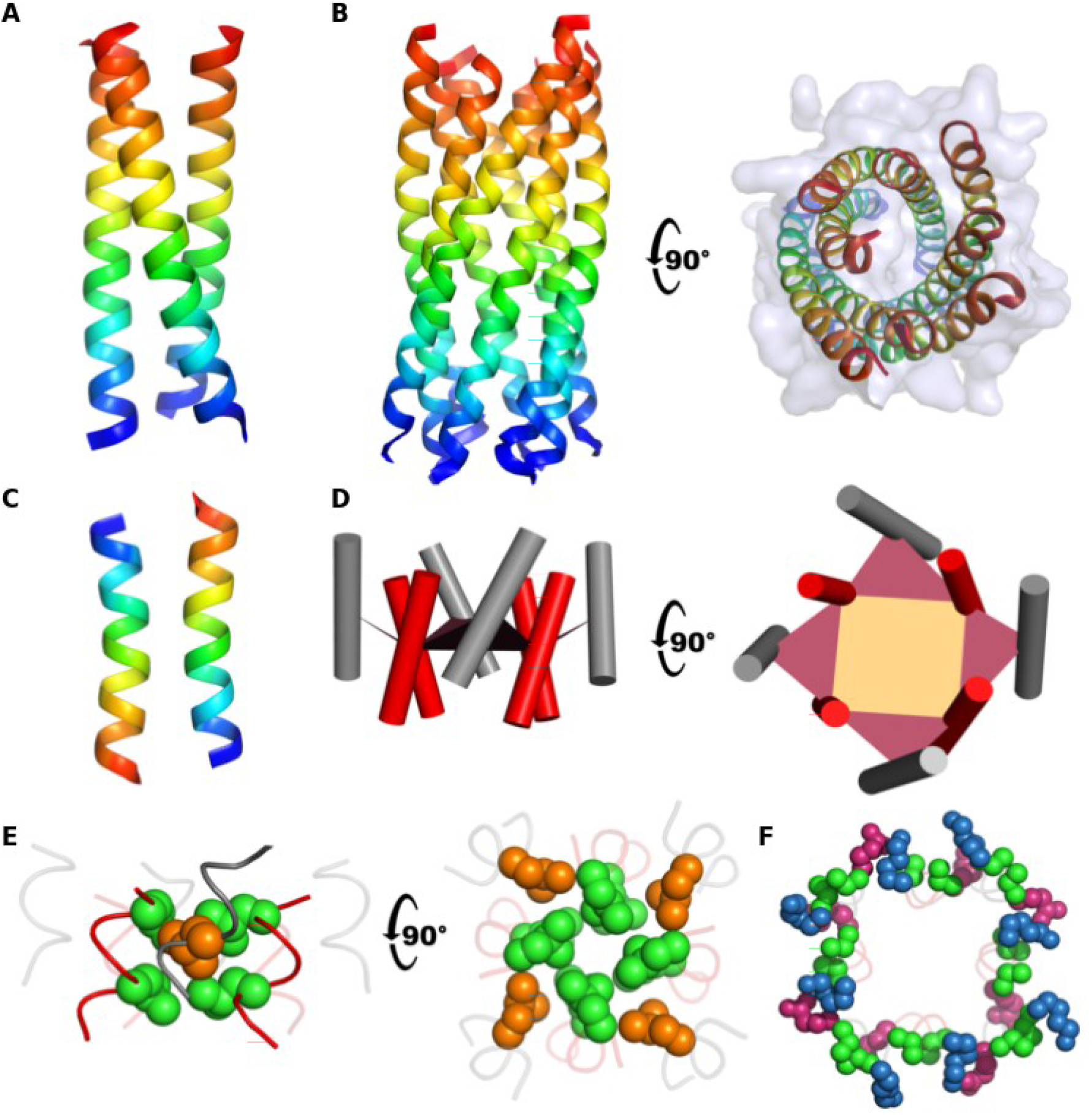
*Crystal structures of de* novo designed peptides. Backbone ribbon structures coloured blue to red from the *N* to *C* termini for: (**A**) PK-1 (CC-TypeN-L_a_L_d_) (PDB id 7qdk); (**B**) orthogonal views of PK-5 (D-3_10_HD) (7qdi); and (**C**) PK-10/PK-11 racemate (7qdj). (**D**) Orthogonal views of PK-5 (D-3_10_HD) showing the outer helices (grey cylinders) shifted with respect to the inner helices (red cylinders), with planes drawn from the centres of identical sequence repeats. (**E**) Similar to panel **D** with Leu side chains shown in space-filling representation to highlight the intimate packing to form a consolidated hydrophobic core. (**F**) Packing of D-Glu (magenta) and D-Lys (blue) residues, plus Aib (green) residues in space-filling representations.

Next, as 3_10_-helix conformations can be stabilised by including Aib residues, we replaced the two Ala residues in each 7-residue repeat of PK-1 to give PK-2, **Table 1**. However, like its parent, PK-2 remained a highly α-helical, thermally stable, and monodisperse trimer in solution, **Fig S3.2** and **S4.2**. Thus, swapping in Aib did not override the propensity of 7-residue repeats to form α-helical bundles. Presumably, this is because the alternating 3,4-pattern of hydrophobic residues, which directs amphipathic α-helix formation and tight coiled-coil packing^37,38^, is not simply overridden by including 3_10_-helix-favouring Aib residues. Therefore, our next step was to break this pattern and find an alternative repeat to favour 3_10_-helix conformations.

On average, the 3_10_ helices of our dataset had 3.28 residues per turn, **Fig. 1C**; though the ideal 3_10_-helix parameter is often stated as 3 residues per turn^10^. Therefore, we reasoned that 3_10_-helical bundles might be the stabilised by reducing the sequence repeat to 3 residues with a single hydrophobic residue per repeat. Our contention was that in an ideal 3_10_ helix, these residues would form a continuous hydrophobic ‘seam’ of an amphipathic 3_10_ helix to promote intermolecular association in water, **Fig. 1G**. Also, by analogy with their use in designed coiled-coil bundles ^35^, we hypothesised that inter-chain salt bridges might stabilise bundles of such helices further. This led us to design two peptides, PK-3 and PK-4 (**Table 1**), with 4 × 6-residue repeats, E-L-Z-Z-L-K, with Z = all Ala (A) or all Aib (U), respectively.

In aqueous buffer, by CD spectroscopy, PK-3 was largely α helical with a broad thermal unfolding transition, **Fig S3.3**. These CD data did not show any dependence on peptide concentration. Consistent with this, AUC experiments confirmed PK-3 as a monodisperse monomer, **Fig S4.3**. The peptide did not crystallise. These data are indicative of single α-helical domains^39^; *i.e*., α-helices that do not associate into higher-order structures. Thus, though the 3-residue hydrophobic repeat removes the drive of the α helix to assemble into bundles, alone it does not stabilise 3_10_ helices or bundles thereof.

By contrast, PK-4 behaved completely differently. First, its CD spectrum was patently different from the three previous peptides, **Fig. 2b**, with a minimum at ≈205 nm characteristic of right-handed helices, but a maximum at ≈220 nm characteristic of right-handed 3_10_ helices in aqueous buffer^40^. The CD spectrum changed little upon heating, **Fig S3.4**. Encouragingly, AUC measurements indicated that PK-4 associated into a monodisperse hexamer, **Fig. 2b** and **Fig S4.4**. *De novo* designed α-helical coiled coils of this size can form barrels^41^ with central lumens that bind the environment-sensitive dye, 1,6-diphenylhexatriene (DPH)^42^. However, DPH did not bind to PK-4 in solution, **Fig S5.1**, suggesting a tightly packed structure with a consolidated hydrophobic core.

We made many attempts to crystallise and solve X-ray structures of PK-4 and its derivatives. This included peptides with L- and D-amino acids, and racemic mixtures of these. Eventually, we crystallised and solved a 2.34 Å resolution structure (**Fig. 3B and Table 7.2&3**) for a variant, PK-5, with D-Glu, D-Lys and D-Leu and a *C*-terminal D-4-bromophenylalanine (D-Br-Phe), **Table 1, Figs. S3.5** and **S4.5**. The structure revealed a parallel bundle of eight left-handed 3_10_ helices with otherwise canonical 3_10_-helix parameters (green asterisks, **Fig. 1B-D**). Notably, the helices have ≈3.15 residues per turn and nine contiguous turns (27 residues) of CO_i_→NH_i+3_ hydrogen bonding. As this is slightly greater than the 3-residue sequence repeat of leucine, these hydrophobic residues track around the left-handed helices in a right-handed manner. This allows the helices to pack in a right-handed supercoil with an extrapolated pitch 93 residues or 177 Å. Long 3_10_ helices in natural proteins tend to be irregular^43^. By contrast, those in the X-ray crystal structure of PK-5 curve smoothly, **Fig S6.1**^44^.

In more detail, the structure of PK-5 has C_4_ symmetry with pairs of adjacent helices contributing to an inner four-helix bundle and an outer four-helix ring, **Fig. 3D**. The packing of the side chains is intimate and reminiscent of ‘knobs-into-holes’ packing in α-helical coiled coils ^37,38^. The Leu residues of the inner helices point directly into a central core, similar to ‘x-layers’ in some α-helical coiled coils ^45^, while those of the outer helices pack into constellations of Leu residues provided by the inner helices, **Fig. 3E**. The latter requires the outer helices to be slipped relative to the inner by ≈2 Å. The solvent-accessible surface of the assembly comprises entirely D-Glu and D-Lys polar residues. In accord with our design strategy, these form a network of salt bridges, with the closest pairs having C_δ_-N_ε_ distances of 3.8 ± 0.5 Å (*N* = 32). Together with the Aib residues, these shield the hydrophobic core from solvent, **Fig. 3F**.

Although both are discrete oligomers, the octameric structure of PK-5 differs in oligomeric state from the hexamer indicated by the solution-phase data of the parent PK-4 (**Fig. 2B**), which has Trp in place of p-Br-Phe. The extended crystal lattice of PK-5 reveals side-by-side and head-to-tail packing of octamers, with the D-p-Br-Phe residues packed in a layer of with many edge-to-face aromatic interactions^46^, **Fig S6.2**. Modelling D-Trp residues into these sites revealed potential steric clashes between the octamer assemblies. Thus, the powerful tendency to assemble programmed into the *de novo* 3_10_-helical sequences can be modulated by subtleties in sequence leading to varying degrees of oligomerisation.

Thus, PK-4 and PK-5 form thermostable, parallel, bundles of six to eight 3_10_ helices. These are unprecedented in both the length of the 3_10_-helices involved and in being the first water-soluble, protein-like supramolecular, or quaternary assemblies of such helices. With our design goal achieved, we renamed PK-4 and PK-5 as 3_10_HD and D-3_10_HD (3_10_-Helical Designs), respectively. This success raises further questions: why are similar structures not found in natural proteins? And, which features of our *de novo* designed sequence make it so disposed to form stable 3_10_-helix-based quaternary structures? To address these questions, we made variants of 3_10_HD.

To start, we tested the effect of hydrophobicity and steric size at the ‘leucine sites’ of the 6-residue repeats on the folding and assembly of 3_10_HD. First, we replaced all of the Leu residues of 3_10_HD with Ala to give PK-6, **Table 1**. CD and AUC measurements showed that this peptide was a partially α-helical (≈50%) monomer in solution, **Fig S3.6** and **S4.6**. Similarly, when the Leu residues were replaced by Aib in PK-7, **Table 1**, the peptide remained monomeric but with an unusual CD spectrum, **Fig S3.7** and **S4.7**. The β-branched hydrophobic residues, Ile and Val, have low propensities for 3_10_ helicity in natural structures, **Fig. 1E**. To examine this, we changed all of Leu residues to Ile (PK-8) or Val (PK-9), **Table 1**. Neither peptide gave a CD spectrum consistent with 3_10_ helicity (**Figs. S3.8** and **S3.9**), and both were monomers in solution (**Figs. S4.8** and **S4.9**). Combined, these experiments indicate that, of the aliphatic hydrophobic side chains, Leu best promotes the supramolecular assembly and stabilisation of 3_10_ helices.

Next, we tested the effect of changing peptide length on the stability of the 3_10_-helix bundle. We kept the overall sequence repeat, ELUULK, but systematically decreased the number of these from 4 to 3 and then to 2 repeats, giving PK-12 and PK-10, respectively, **Table 1**. The shortest peptide, PK-10, was a partly folded monomer in solution, **Figs. S3.10** and **S4.10**; and a structure of a racemic mixture (PK-10+PK-11) at 1.4 Å resolution, **Fig. 3C**, revealed two antiparallel α helices packed with ‘knobs-into-holes’ interactions as observed in other heterochiral systems^38,47^. By contrast, and interestingly, CD spectra of PK-12 unveiled a concentration-dependent switch from a partially α-helical to a 3_10_-helical conformation, **Fig. 2C** and **Fig. S3.11**. As the peptide concentration was increased from 5 μM to 1 mM, the negative maximum at 226 nm changed to a positive maximum at 220 nm. Concomitantly, the oligomeric state of the peptide changed from multiple low-order species at 25 μM to hexamer to hexamer at higher concentrations, **Figs. S4.11&12**. Attempts to crystallise PK-12 failed. Nonetheless, the series PK-4 (3_10_HD)→PK-12→PK-10 shows that peptide length, and with it the length of the hydrophobic seam, is critical for the folding, assembly, and stabilisation of 3_10_ helices.

Finally, exploiting the symmetry and salt bridging of the D-3_10_HD structure, we designed a heteromeric system comprising acidic, PK-13, and basic, PK-14, peptides, **Table 1**. In isolation in solution, the two peptides were partially folded, α-helical monomers, **Fig. 2D (top)** and **Figs. S4.13&14**. However, when mixed, the CD spectrum switched to that of the parent homomeric assembly, **Figs. 2B&D**, indicating 3_10_-helix formation. Moreover, a Job plot gave a 1:1 stoichiometry for the complex formed, **Fig. 2D (middle)** and **S3.12**. Finally, AUC measurements returned a monodisperse hexamer similar to the parent peptide, 3_10_HD, **Figs. 2B&D (bottom)** and **Figs. S4.4&15**. Retrospectively, we named this completely new *de novo* designed heteromeric quaternary complex of 3_10_ helices 3_10_HD-AB.

In conclusion, we have achieved the rational *de novo* design of unprecedented water-soluble supramolecular assemblies, or protein-like quaternary structures, constructed from 3_10_-helical peptides. The designs incorporate: (i) a bioinformatically guided reduced amino-acid alphabet; (ii) the non-proteinogenic α,α-disubstituted amino acid, Aib; (iii) strict 6-residue sequence repeats; and (iv) a minimum number of 3 such repeats. All but one of these features (*i.e*., number ii) could be achieved through natural ribosomal protein synthesis, which raises the question: why has nature not found and exploited these or similar structures? The significance of our successful designs is that they require extended and amphipathic 3_10_ helices, which are rare in nature. We suggest that these requirements are key, otherwise polypeptide chains will adopt alternative α-helical conformations, which are nearby in conformational space and energetically more favourable. Nonetheless, we provide decisive proof that this can be overridden to achieve stabilised 3_10_-helical conformations and assemblies. It will be interesting to see what explorations of new structural and functional peptide chemistry this route into a seemingly unexplored region of protein-structure space opens up. For instance, Aib and similar residues can be incorporated into engineered proteins^48^, the chemistry of α,α-disubstituted amino acids is being expanded^49^, and other peptide and related foldamers are increasingly being exploited as functional frameworks for binding and catalysis^50,51^.

## Acknowledgments

PK and DNW are supported by Biotechnology and Biological Sciences Research Council (BBSRC) grant to DNW (BB/R00661X/1). DNW is also supported by BrisSynBio, a BBSRC/EPSRC-funded Synthetic Biology Research Centre (BB/L01386X/1), and a Royal Society Wolfson Research Merit Award (WM140008). JC is supported by ERC Advanced Grant DOGMATRON (Agreement no. 884786), and by an EPSRC Programme Grant (EP/P027067/1). We thank the University of Bristol School of Chemistry Mass Spectrometry Facility for access to the EPSRC-funded Bruker Ultraflex MALDI-TOF/TOF instrument (EP/ K03927X/1), and BrisSynBio for access to peptide synthesizers. We thank Diamond Light Source for access to beamlines I03, I04, I04-1 and I24 (Proposal 23269) and Dr Mark Warren from I19 who helped NGP with the direct methods solution. The authors thank Prof Todd Yeates (UCLA), Kapil Gupta, Christine Tölzer, Frank Zieleniewski, and members of the Clayden and Woolfson labs and BrisSynBio for helpful discussions.

## Author Contributions

PK, JC and DNW conceived the project. PK and DNW designed the bioinformatics analyses, which were performed by PK. PK and DNW designed the sequences, which were synthesized, characterised, and crystallised by PK. PK and NGP solved the X-ray crystal structures. PK, JC and DNW wrote the manuscript, which was read by all authors.

