## Supplemental Information for "Constructing synthetic-protein assemblies from *de novo* designed 3_10_ helices"

| | |
|---|---|
| Section 1 | Bioinformatics Analyses |
| Section 2 | Analytical High-Pressure Liquid Chromatography (HPLC) and Matrix-Assisted Laser Desorption/Ionisation - Time of Flight (MALDI-TOF) |
| Section 3 | Circular Dichroism (CD) Spectroscopy |
| Section 4 | Analytical Ultracentrifugation (AUC) |
| Section 5 | DPH-binding Analyses |
| Section 6 | Structural Analyses of $3_{10}$-helix bundle |
| Section 7 | Tables |

**Section 1: Bioinformatics Analyses**

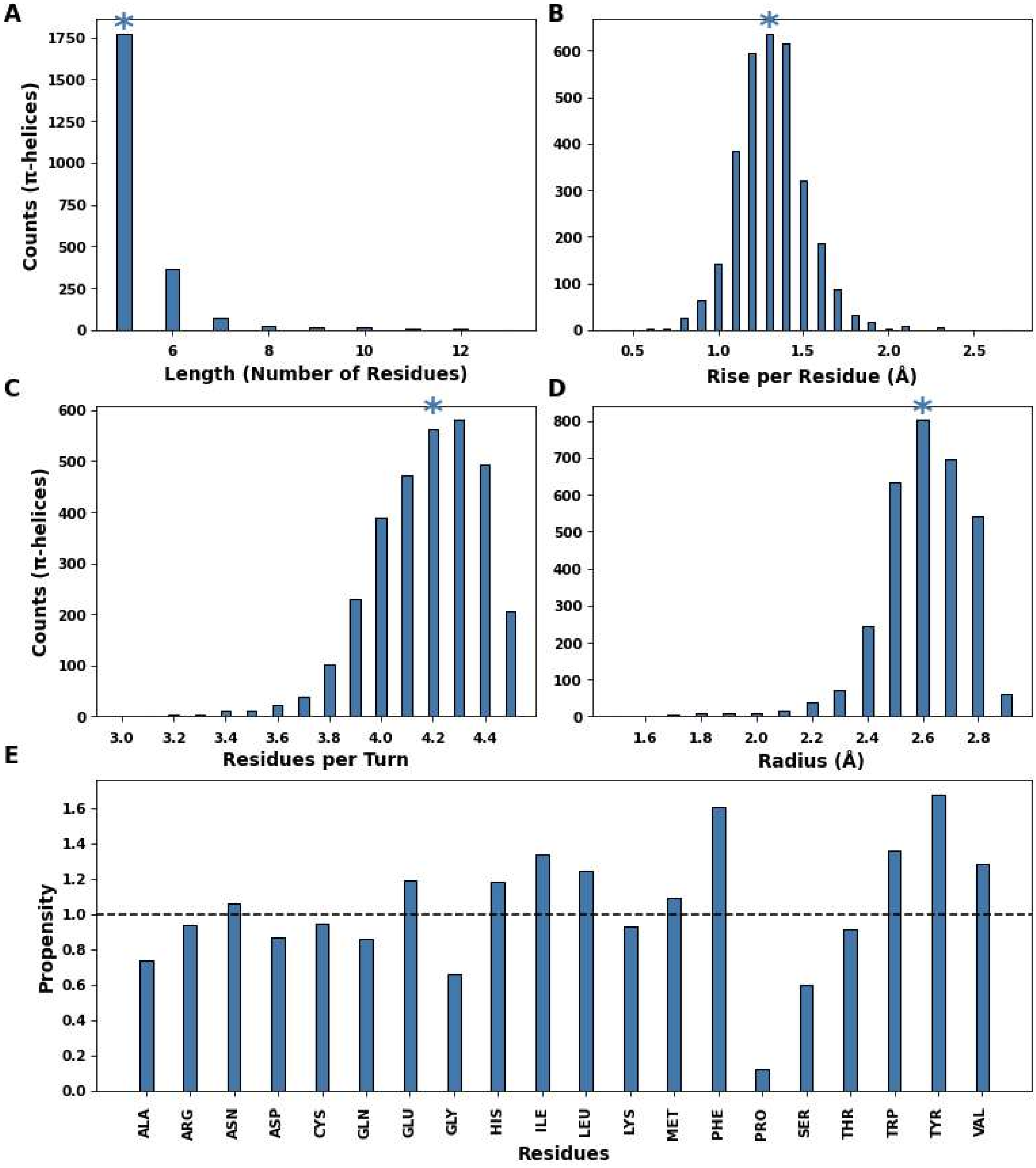


**Figure S1.1: Analysis of the π helices as identified by DSSP in the subset of the PDB.** Helical parameters of π helices: (**A**) helix length; (**B**) rise per residue; (**C**) residues per turn; and (**D**) radius. Mean values are marked with blue asterisks. (**E**) Propensity values of the 20 standard amino acids in π helices.

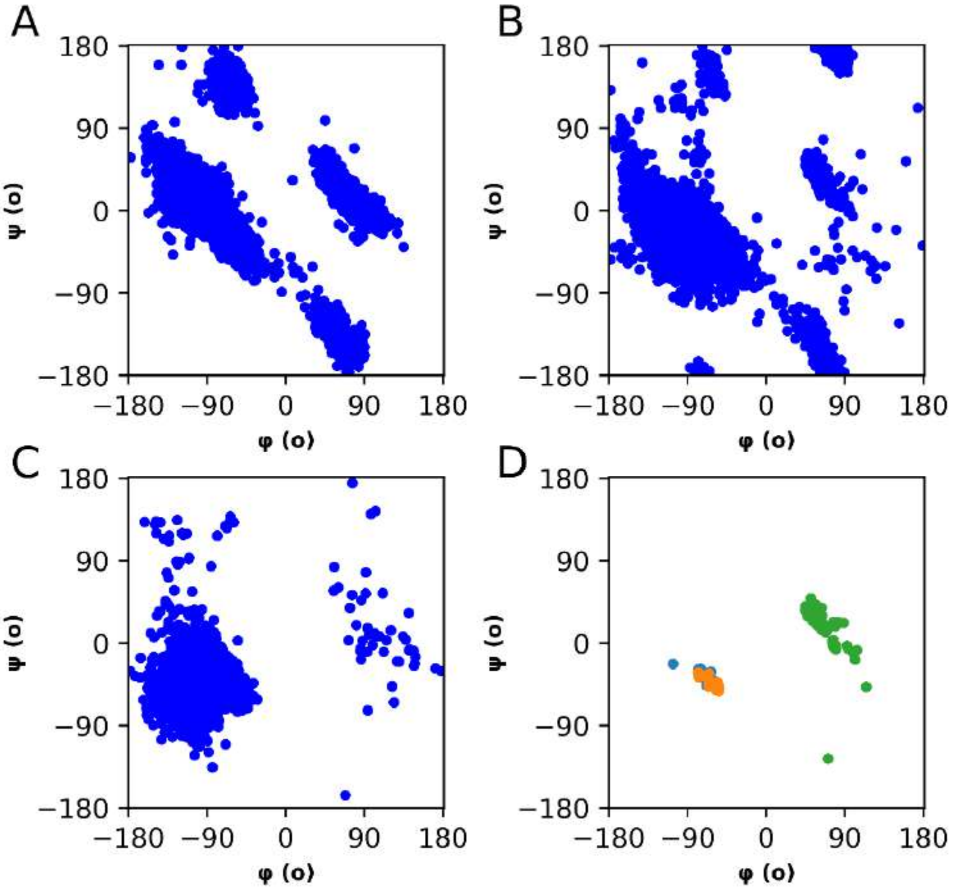


**Figure S1.2: Distributions of backbone Ramachandran angles.** (**A**) For 90,281 residues in 26,917 $3_{10}$ helices from the subset of the PDB; (**B**) 742,703 residues in 66,771 α helices; and (**C**) 12,196 residues in 2,294 π helices. Calculated mean and standard deviations are listed in **Table S7.1**. (**D**) For the X-ray crystal structures determined herein: CC-TypeN-$L_aL_d$ (PK-1, orange); D-$3_{10}$-HD (PK-5, green); and PK-10 (blue).

**Section 2: Matrix-Assisted Laser Desorption/Ionisation - Time of Flight (MALDI-TOF) and Analytical High-Pressure Liquid Chromatography (HPLC)**

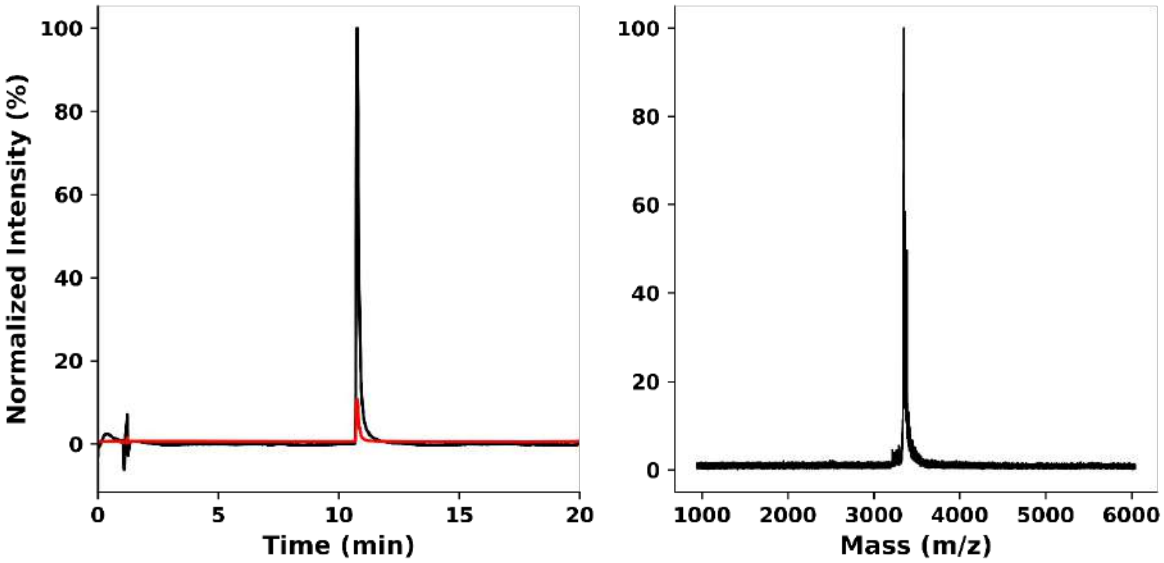


**Figure S2.1:** Characterisation of PK-1 (CC-TypeN-$L_aL_d$). **Left:** Analytical HPLC chromatogram monitored at 220 nm (red) and 280 nm (black). **Right**: MALDI-TOF MS. Calculated mass: 3345.8 Da. Observed mass: 3347.0 Da $[M+H]^+$.

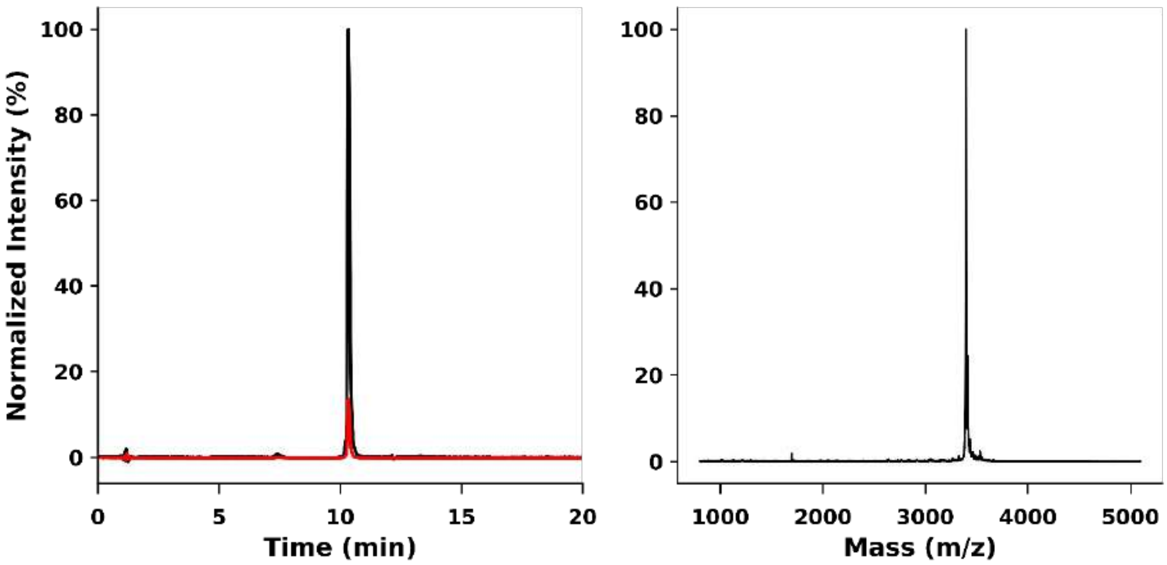


**Figure S2.2:** Characterisation of PK-2. **Left:** Analytical HPLC chromatogram monitored at 220 nm (red) and 280 nm (black). **Right**: MALDI-TOF MS. Calculated mass: 3398.1 Da. Observed mass: 3398.1 Da.

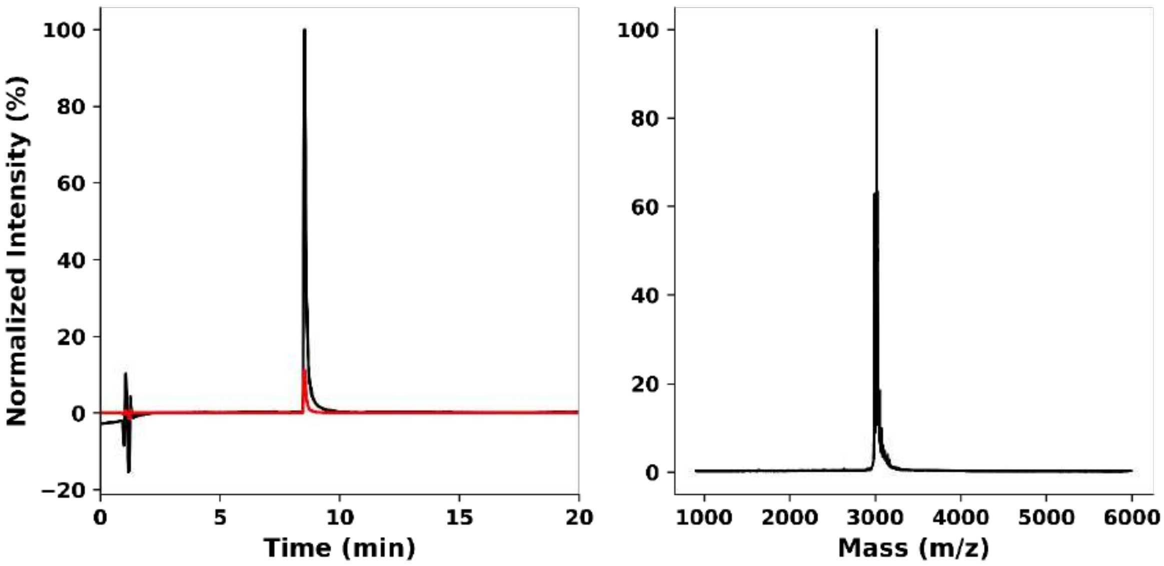


**Figure S2.3:** Characterisation of PK-3. **Left:** Analytical HPLC chromatogram monitored at 220 nm (red) and 280 nm (black). **Right**: MALDI-TOF MS. Calculated mass: 2990.6 Da. Observed mass: 3014.1 Da $[M+Na]^+$.

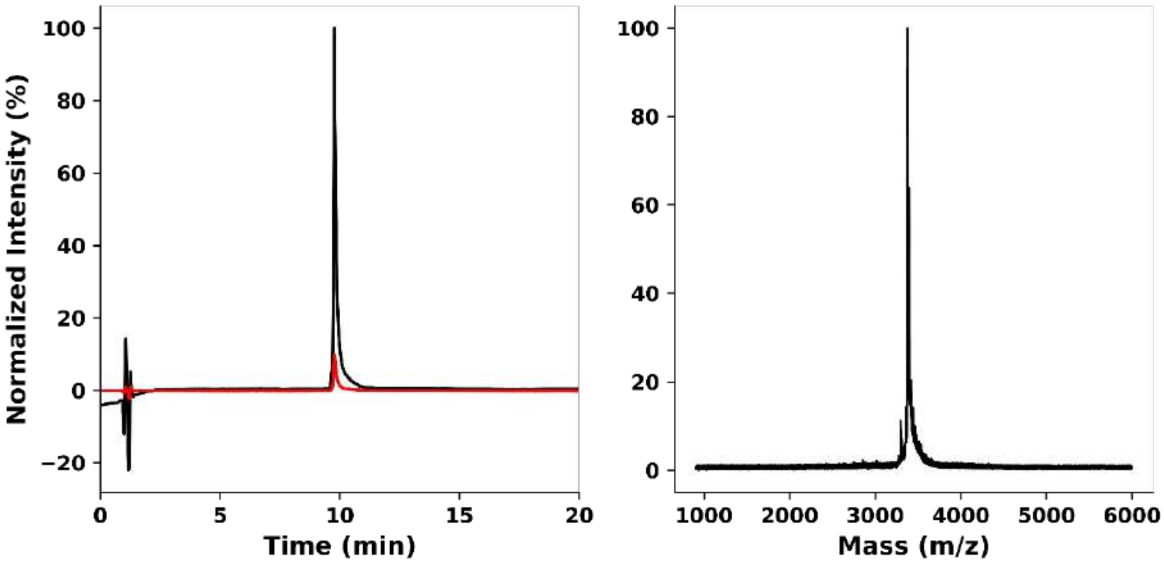


**Figure S2.4:** Characterisation of PK-4 ($3_{10}$HD). **Left:** Analytical HPLC chromatogram monitored at 220 nm (red) and 280 nm (black). **Right**: MALDI-TOF MS. Calculated mass: 3358.1 Da. Observed mass: 3380.7 Da $[M+Na]^+$.

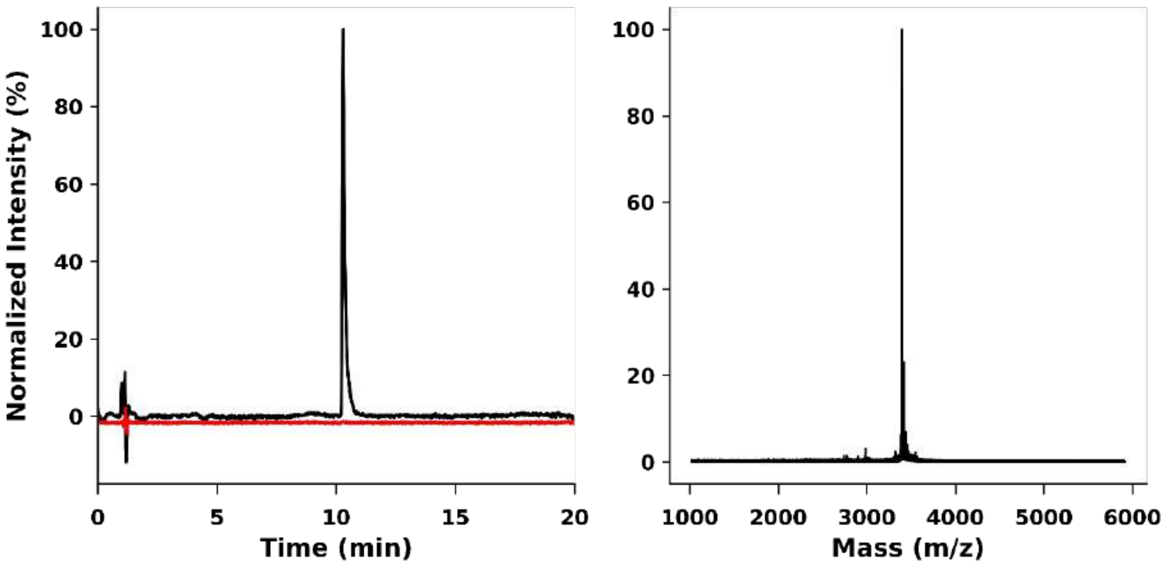


**Figure S2.5:** Characterisation of PK-5 (D-$3_{10}$HD). **Left:** Analytical HPLC chromatogram monitored at 220 nm (red) and 280 nm (black). **Right**: MALDI-TOF MS. Calculated mass: 3398.1 Da. Observed mass: 3398.1 Da $[M+H]^+$.

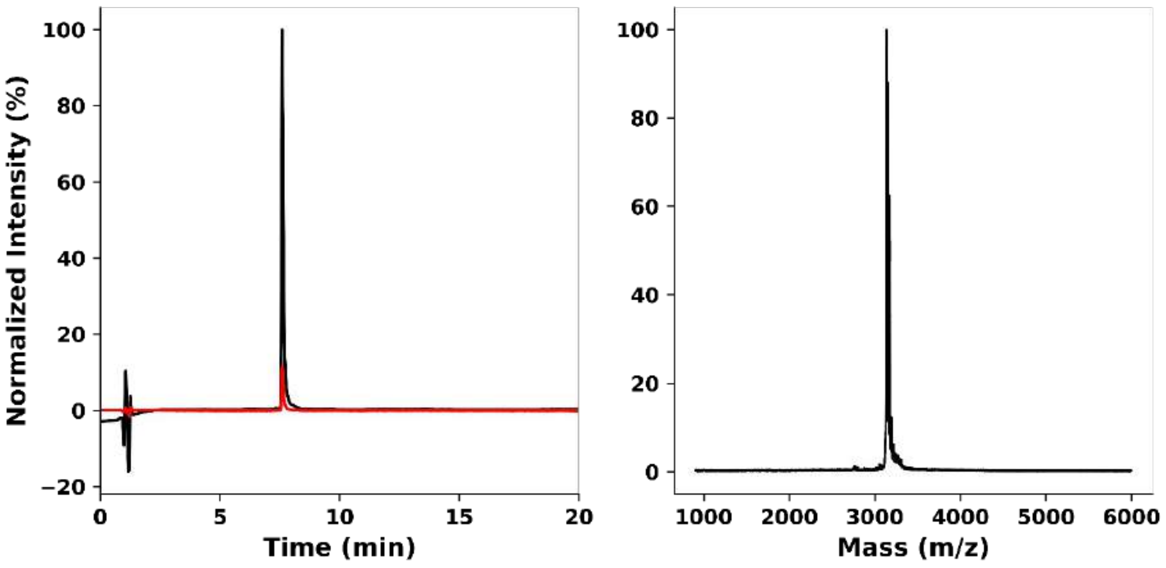


**Figure S2.6:** Characterisation of PK-6. **Left:** Analytical HPLC chromatogram monitored at 220 nm (red) and 280 nm (black). **Right**: MALDI-TOF MS. Calculated mass: 3021.5 Da. Observed mass: 3022.1 Da $[M+H]^+$.

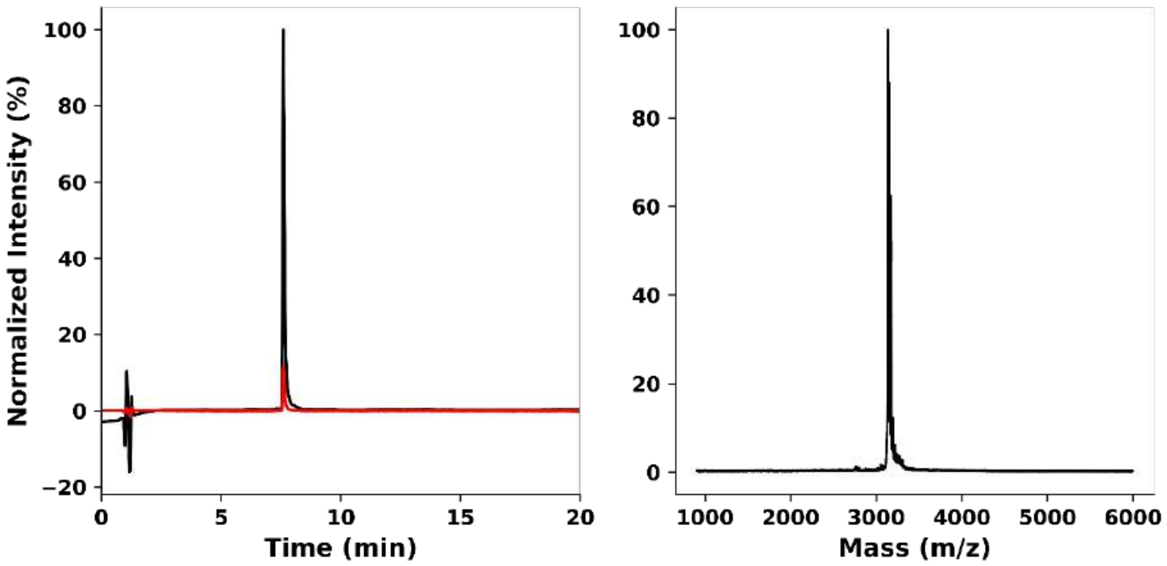


**Figure S2.7:** Characterisation of PK-7. **Left:** Analytical HPLC chromatogram monitored at 220 nm (red) and 280 nm (black). **Right**: MALDI-TOF MS. Calculated mass: 3133.7 Da. Observed mass: 3134.9 Da $[M+H]^+$.

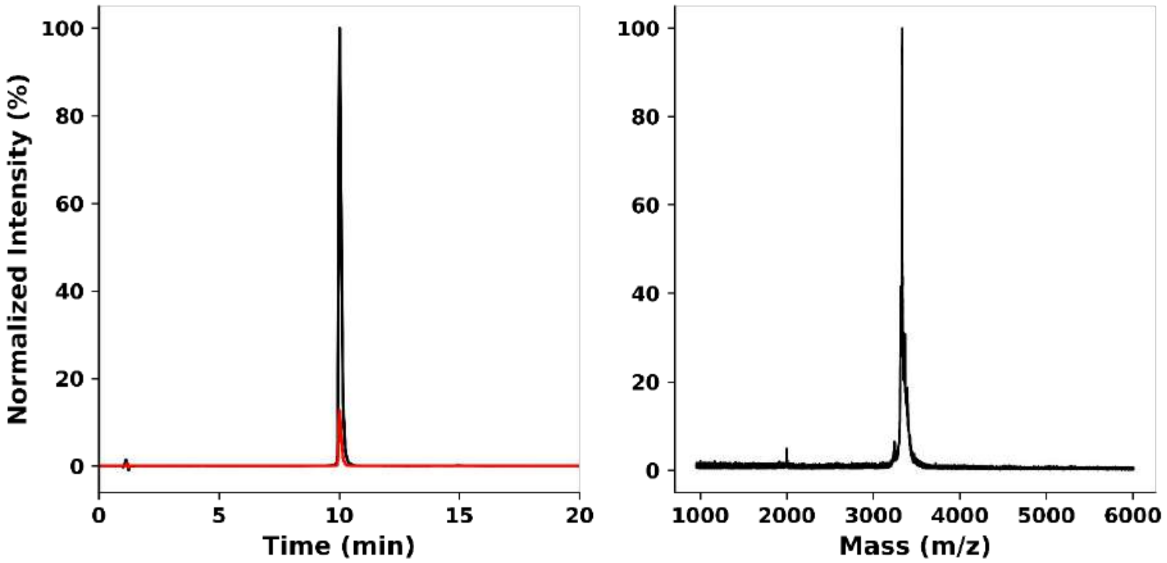


**Figure S2.8:** Characterisation of PK-8. **Left:** Analytical HPLC chromatogram monitored at 220 nm (red) and 280 nm (black). **Right**: MALDI-TOF MS. Calculated mass: 3356.0 Da. Observed mass: 3357.0 Da $[M+H]^+$.

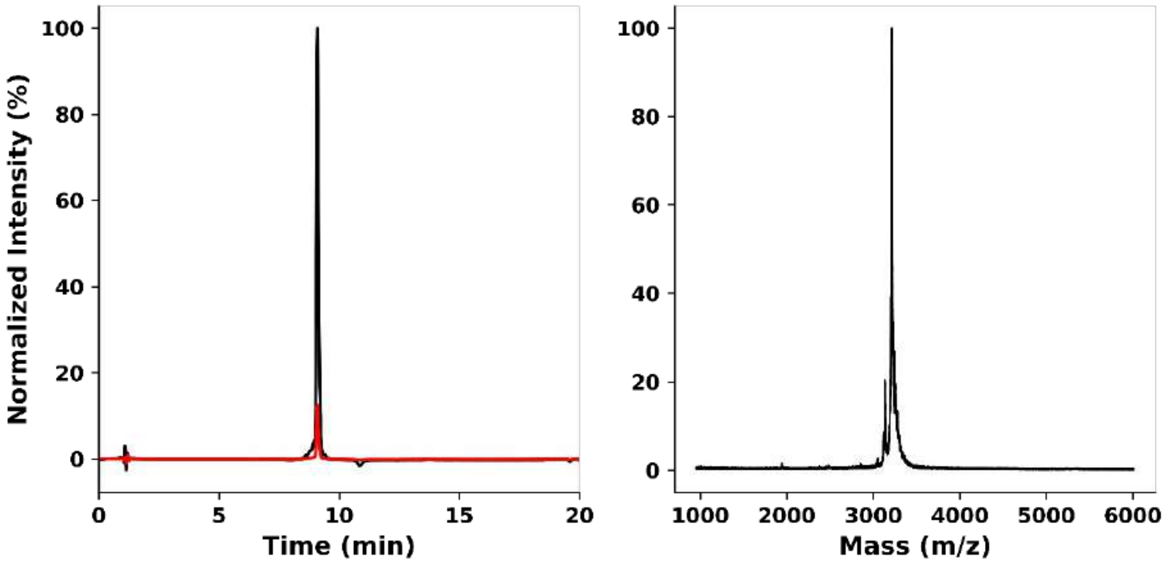


**Figure S2.9:** Characterisation of PK-9. **Left:** Analytical HPLC chromatogram monitored at 220 nm (red) and 280 nm (black). **Right**: MALDI-TOF MS. Calculated mass: 3243.9 Da. Observed mass: 3243.9 Da.

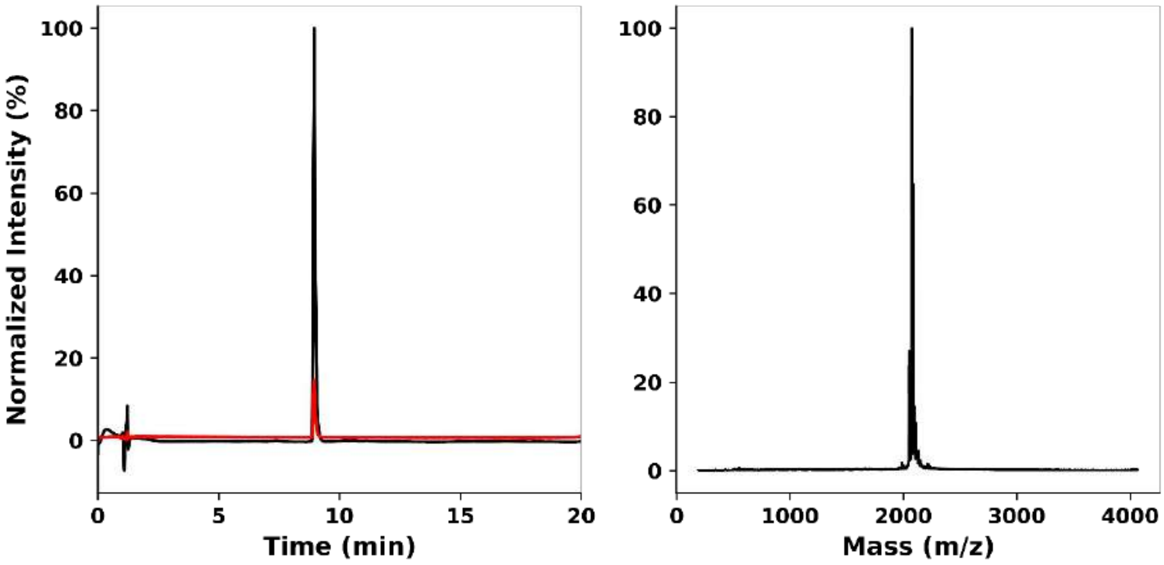


**Figure S2.10:** Characterisation of PK-10. **Left:** Analytical HPLC chromatogram monitored at 220 nm (red) and 280 nm (black). **Right**: MALDI-TOF MS. Calculated mass: 2051.4 Da $[M+H]^+$. Observed mass: 2074 Da $[M+Na]^+$.

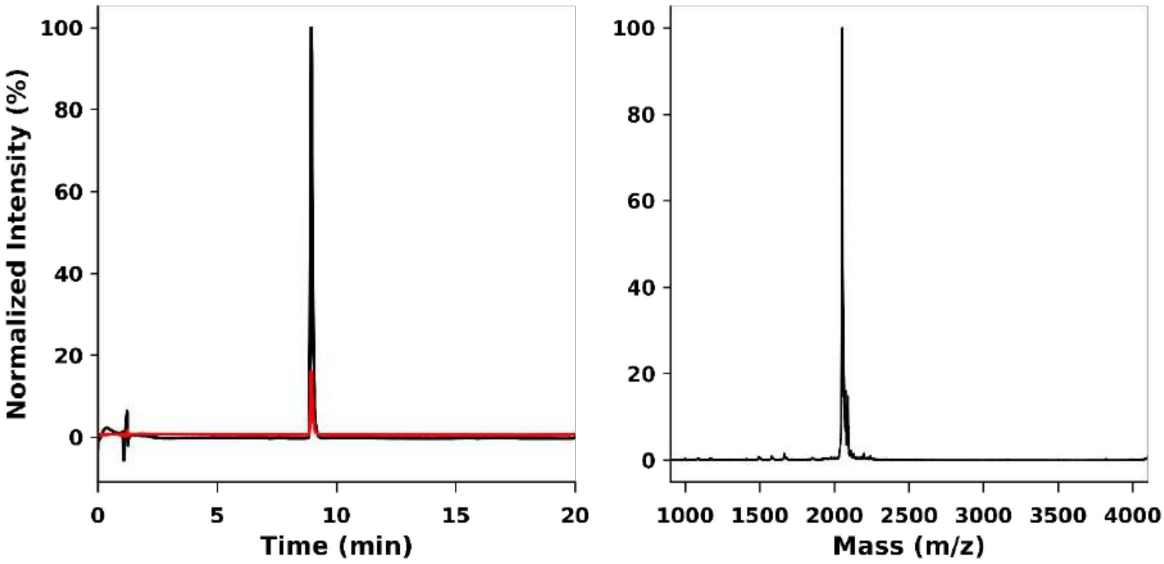


**Figure S2.11:** Characterisation of PK-11. **Left:** Analytical HPLC chromatogram monitored at 220 nm (red) and 280 nm (black). **Right**: MALDI-TOF MS. Calculated mass: 2051.4 Da. Observed mass: 2053.0 Da $[M+H]^+$.

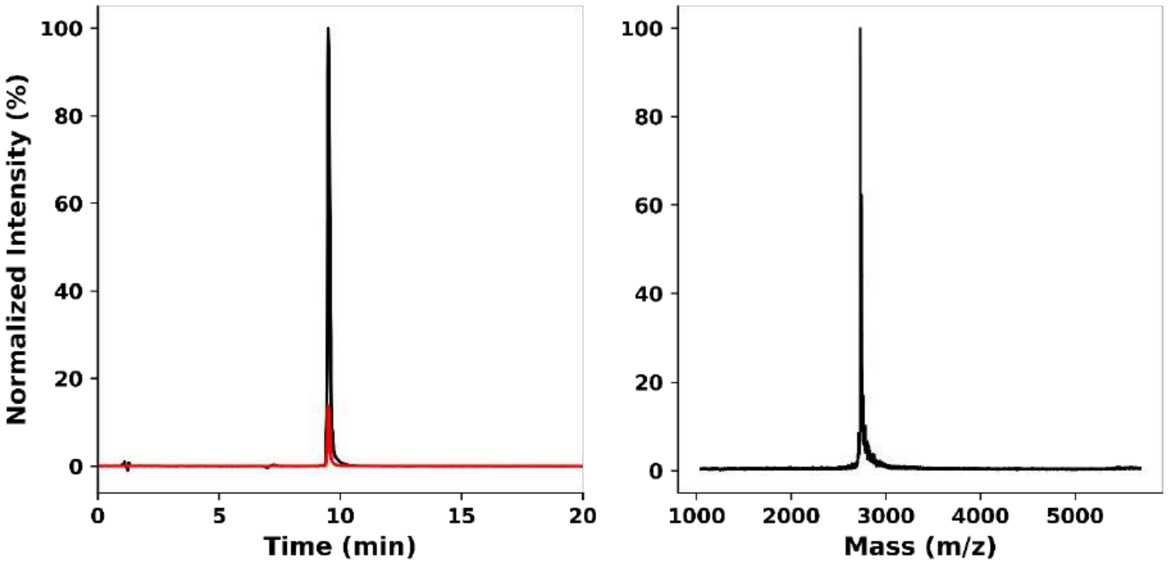


**Figure S2.12:** Characterisation of PK-12. **Left:** Analytical HPLC chromatogram monitored at 220 nm (red) and 280 nm (black). **Right**: MALDI-TOF MS. Calculated mass: 2704.3 Da. Observed mass: 2728.3 Da $[M+Na]^+$.

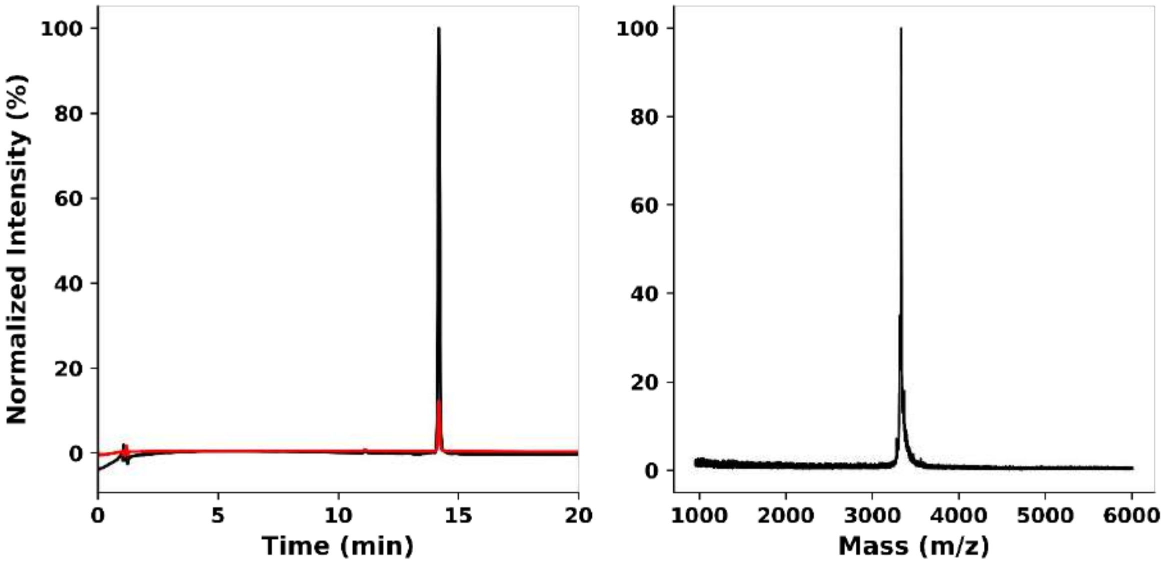


**Figure S2.13:** Characterisation of PK-13 ($3_{10}$HD-A). **Left:** Analytical HPLC chromatogram monitored at 220 nm (red) and 280 nm (black). **Right**: MALDI-TOF MS. Calculated mass: 3359.8 Da. Observed mass: 3359.7 Da.

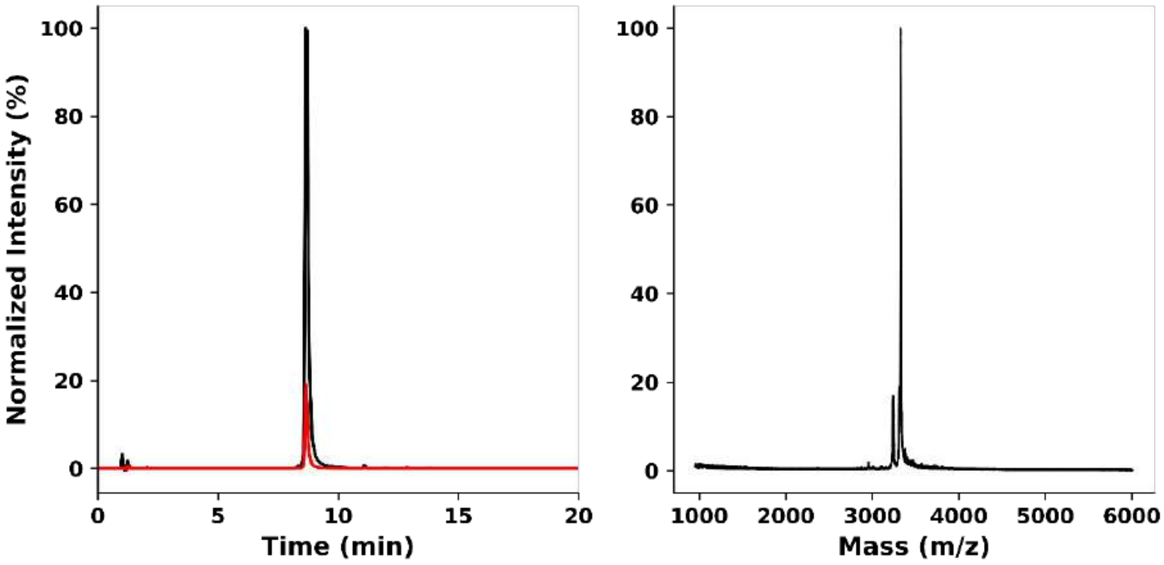


**Figure S2.14:** Characterisation of PK-14 ($3_{10}$HD-B). **Left:** Analytical HPLC chromatogram monitored at 220 nm (red) and 280 nm (black). **Right**: MALDI-TOF MS. Calculated mass: 3352.3 Da. Observed mass: 3353.0 Da $[M+H]^+$.

**Section 3 Circular Dichroism (CD)**

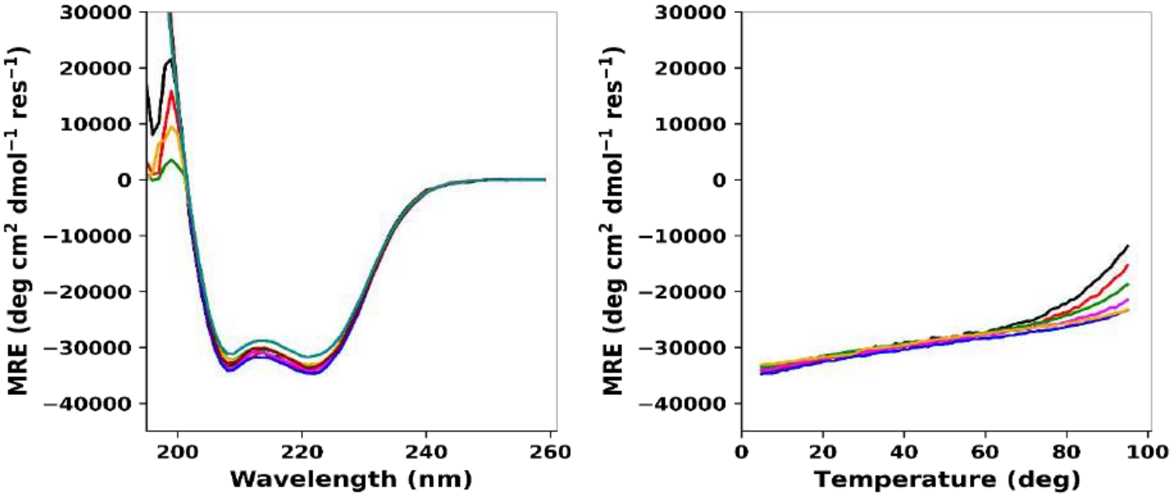


**Figure S3.1:** CD spectra of PK-1 (CC-TypeN-$L_aL_d$) at 5 °C (**left**) and thermal denaturation profile monitored at 222 nm (**right**). Peptide concentrations: 5 µM (black), 10 µM (red), 25 µM (green), 50 µM (magenta), 100 µM (blue), 200 µM (orange), 500 µM (maroon), 1000 µM (light green). Conditions: PBS (pH 7.4). Thermal denaturation profile for peptide concentrations 500 µM and 1000 µM are not plotted.

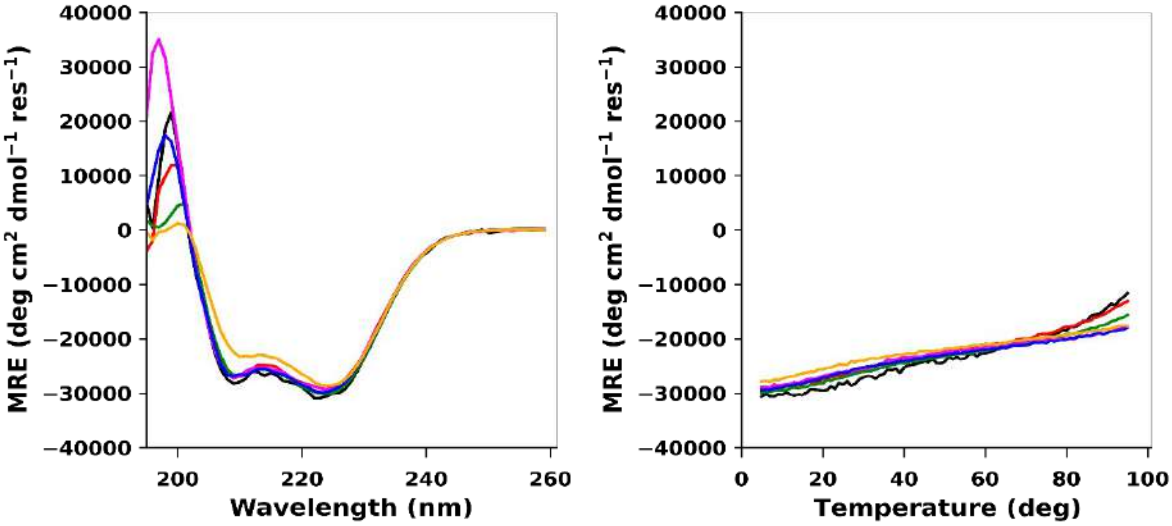


**Figure S3.2:** CD spectra of PK-2 at 5 °C (**left**) and thermal denaturation profile monitored at 222 nm (**right**). Peptide concentrations: 5 µM (black), 10 µM (red), 25 µM (green), 50 µM (magenta), 100 µM (blue) and 200 µM (orange). Conditions: PBS (pH 7.4). Peptide was insoluble after pH adjustment at concentrations 500 µM and 1000 µM.

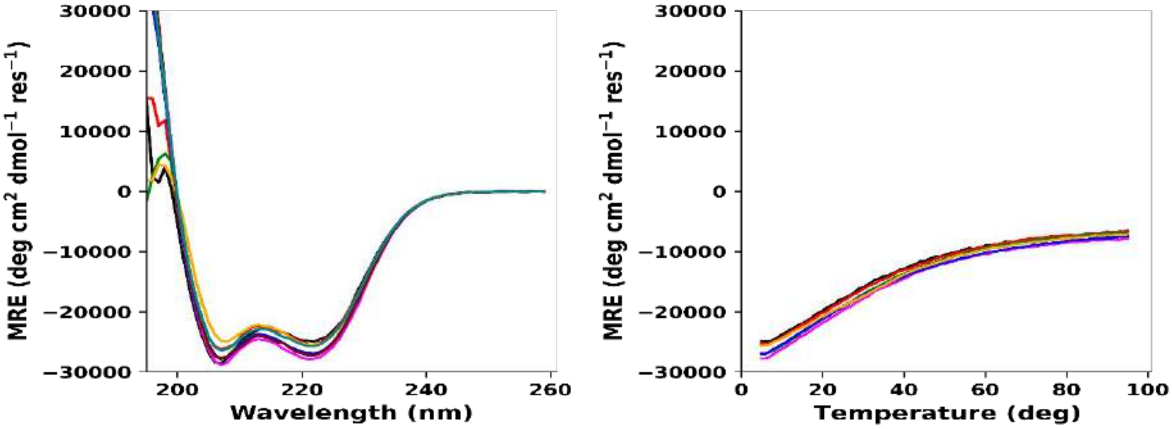


**Figure S3.3:** CD spectra of PK-3 at 5 °C (**left**) and thermal denaturation profile monitored at 222 nm (**right**). Peptide concentrations: 5 µM (black), 10 µM (red), 25 µM (green), 50 µM (magenta), 100 µM (blue), 200 µM (orange), 500 µM (maroon), 1000 µM (light green). Conditions: PBS (pH 7.4). Thermal denaturation profile for peptide concentrations 500 µM and 1000 µM are not plotted.

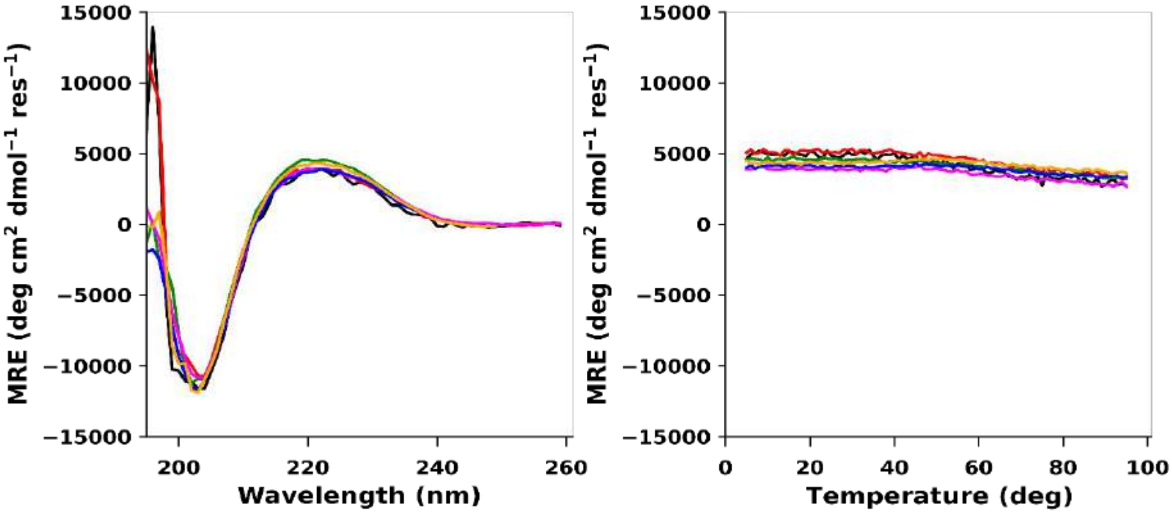


**Figure S3.4:** CD spectra of PK-4 ($3_{10}$HD) at 5 °C (**left**) and thermal denaturation profile monitored at 222 nm (**right**). Peptide concentrations: 5 µM (black), 10 µM (red), 25 µM (green), 50 µM (magenta), 100 µM (blue), 200 µM (orange), 500 µM (maroon), 1000 µM (light green). Conditions: PBS (pH 7.4). Thermal denaturation profile for peptide concentrations 500 µM and 1000 µM are not plotted.

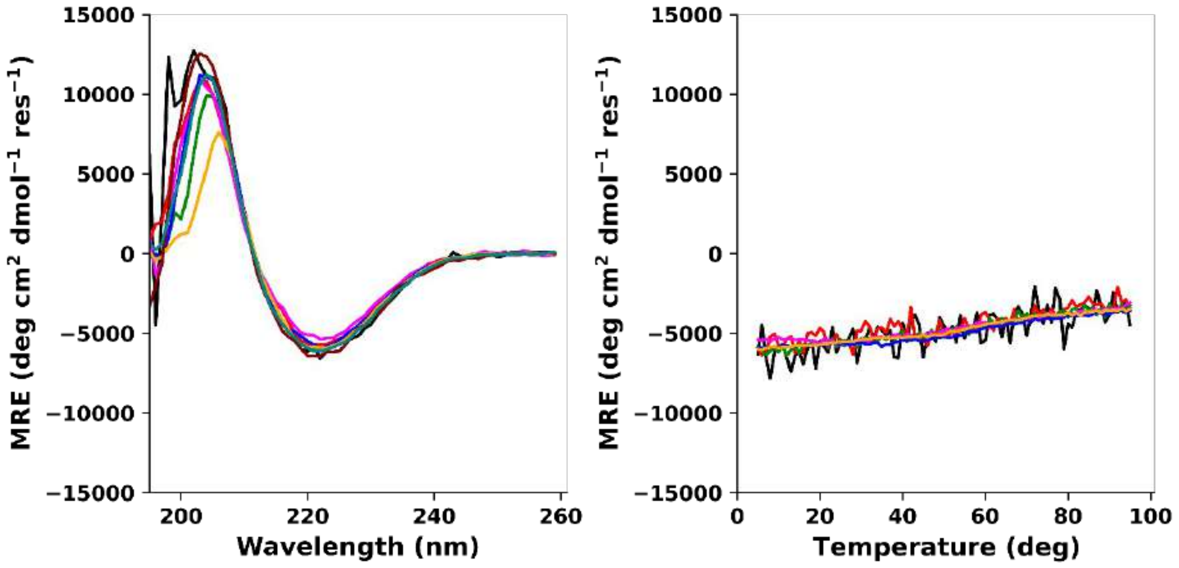


**Figure S3.5:** CD spectra of PK-5 (D-$3_{10}$HD) at 5 °C (**left**) and thermal denaturation profile monitored at 222 nm (**right**). Peptide concentrations: 5 µM (black), 10 µM (red), 25 µM (green), 50 µM (magenta), 100 µM (blue), 200 µM (orange), 500 µM (maroon), 1000 µM (light green). Conditions: PBS (pH 7.4). Thermal denaturation profile for peptide concentrations 500 µM and 1000 µM are not plotted.

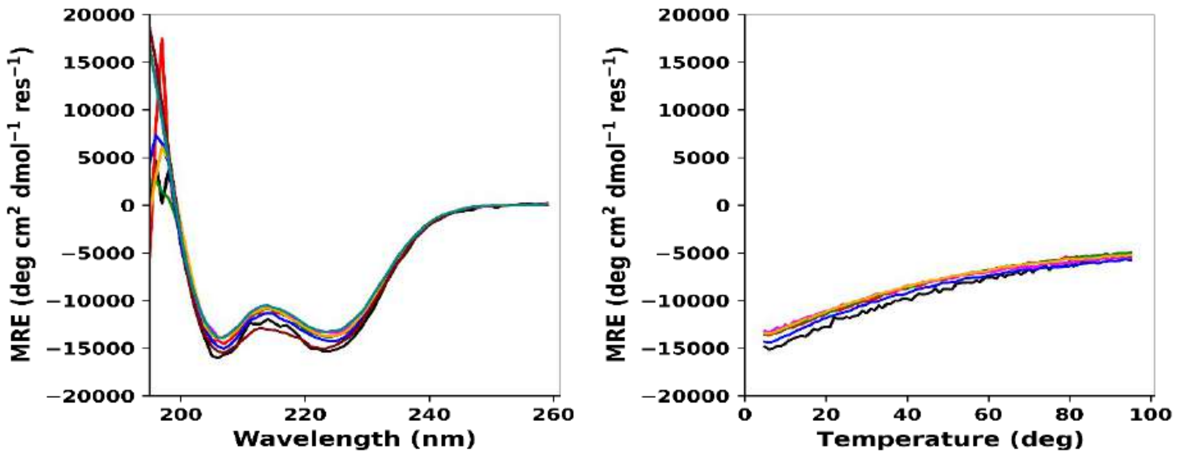


**Figure S3.6:** CD spectra of PK-6 at 5 °C (**left**) and thermal denaturation profile monitored at 222 nm (**right**). Peptide concentrations: 5 µM (black), 10 µM (red), 25 µM (green), 50 µM (magenta), 100 µM (blue), 200 µM (orange), 500 µM (maroon), 1000 µM (light green). Conditions: PBS (pH 7.4). Thermal denaturation profile for peptide concentrations 500 µM and 1000 µM are not plotted.

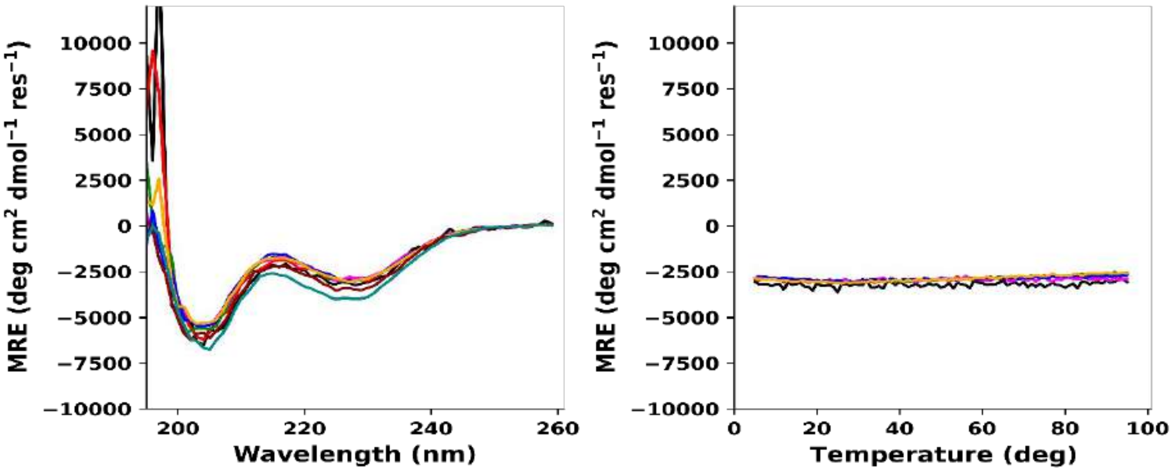


**Figure S3.7:** CD spectra of PK-7 at 5 °C (**left**) and thermal denaturation profile monitored at 222 nm (**right**). Peptide concentrations: 5 µM (black), 10 µM (red), 25 µM (green), 50 µM (magenta), 100 µM (blue), 200 µM (orange), 500 µM (maroon), 1000 µM (light green). Conditions: PBS (pH 7.4). Thermal denaturation profile for peptide concentrations 500 µM and 1000 µM are not plotted.

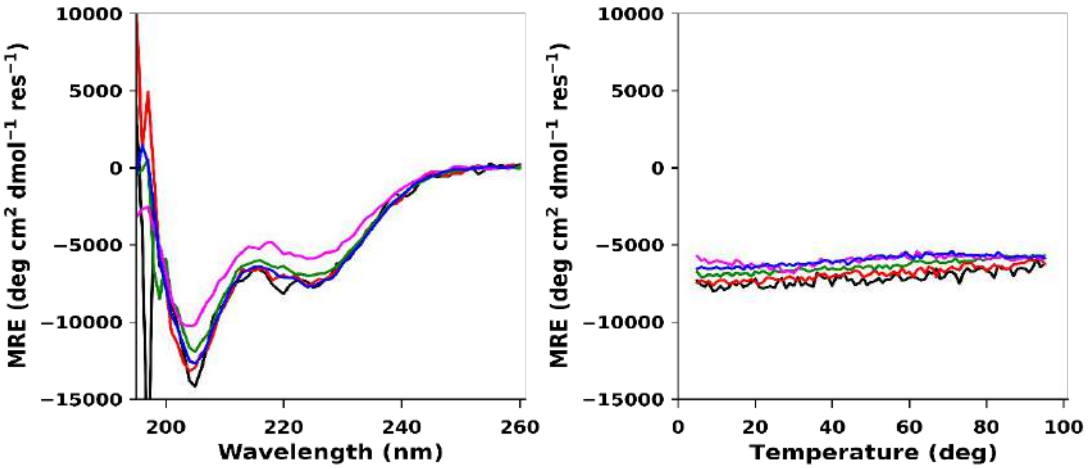


**Figure S3.8:** CD spectra of PK-8 at 5 °C (**left**) and thermal denaturation profile monitored at 222 nm (**right**). Peptide concentrations: 5 µM (black), 10 µM (red), 25 µM (green), 50 µM (magenta), 100 µM (blue). Conditions: PBS (pH 7.4).

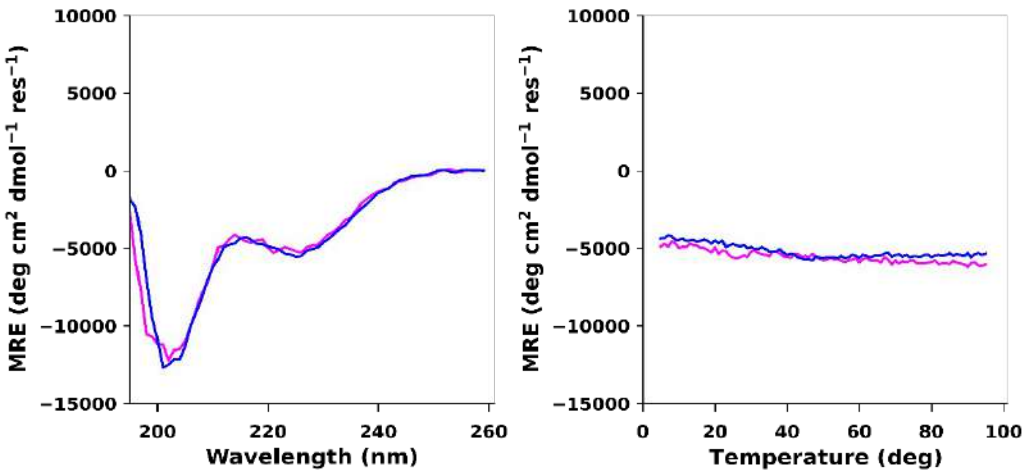


**Figure S3.9:** CD spectra of PK-9 at 5 °C (**left**) and thermal denaturation profile monitored at 222 nm (**right**). Peptide concentrations: 50 µM (magenta), 100 µM (blue). Conditions: PBS (pH 7.4).

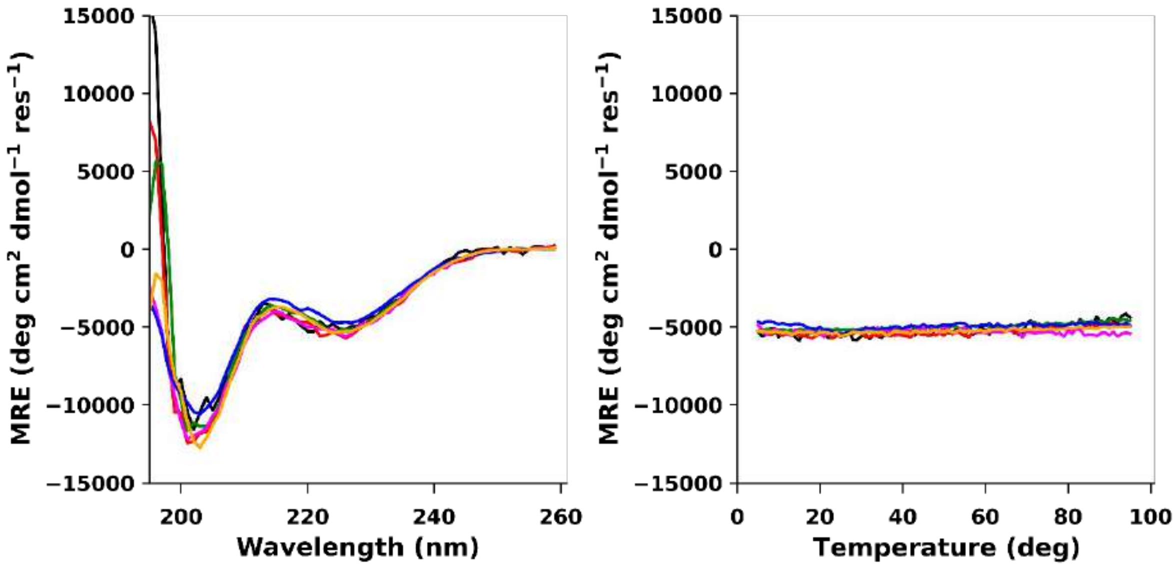


**Figure S3.10:** CD spectra of PK-10 at 5 °C (**left**) and thermal denaturation profile monitored at 222 nm (**right**). Peptide concentrations: 5 µM (black), 10 µM (red), 25 µM (green), 50 µM (magenta), 100 µM (blue), 200 µM (orange), 500 µM (maroon), 1000 µM (light green). Conditions: PBS (pH 7.4). Thermal denaturation profile for peptide concentrations 500 µM and 1000 µM are not plotted.

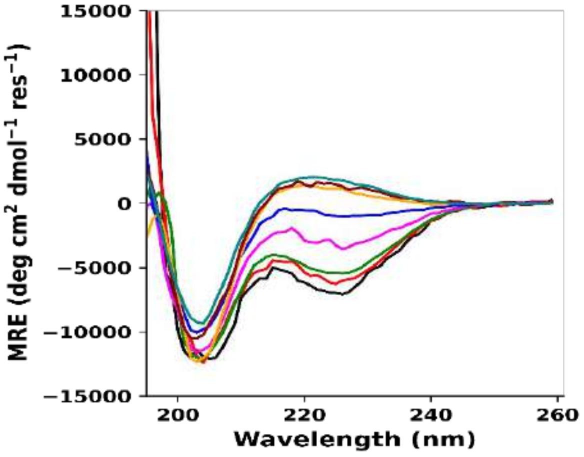


**Figure S3.11:** CD spectra of PK-12 at 5 °C (**left**) and thermal denaturation profile monitored at 222 nm (**right**). Peptide concentrations: 5 µM (black), 10 µM (red), 25 µM (green), 50 µM (magenta), 100 µM (blue), 200 µM (orange), 500 µM (maroon), 1000 µM (light green). Conditions: PBS (pH 7.4). Thermal denaturation profile for peptide concentrations 500 µM and 1000 µM are not plotted.

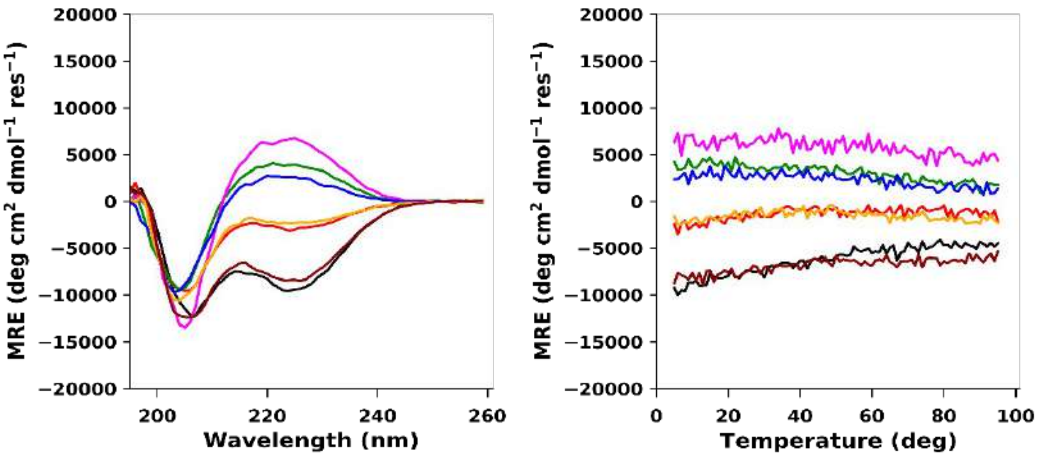


**Figure S3.12:** CD spectra for the mixture of PK-13 ($3_{10}$HD-A) and PK-14 ($3_{10}$HD-B) at a total concentration 100 µM and 5 °C (**left**), and thermal denaturation profile monitored at 222 nm (**right**). Peptide compositions: A100:0B (black), A80:20B (red), A60:40B (green), A50:50B (magenta), A40:60B (blue), A20:80B (orange), A0: 100B (maroon). Conditions: PBS (pH 7.4).

**Section 6: Analytical Ultracentrifugation (SV and SE)**

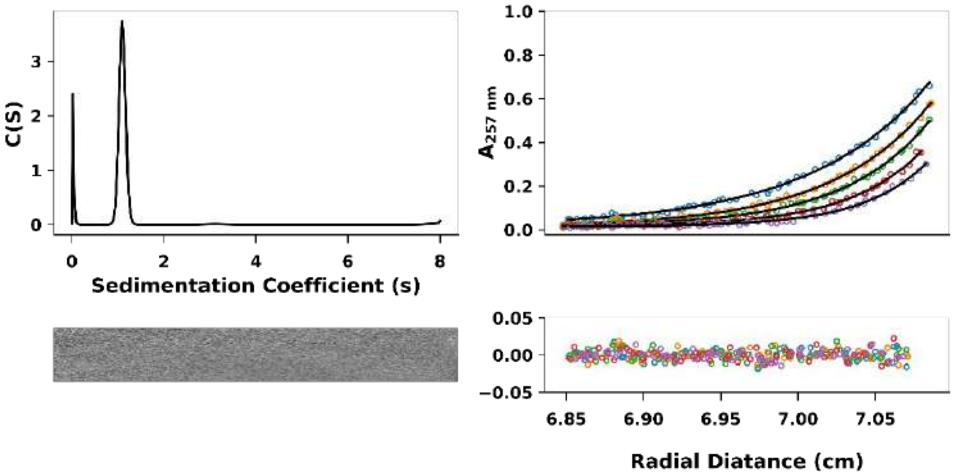


**Figure S4.1:** AUC data and fits (top) with residuals (bottom) for PK-1 (CC-TypeN-$L_aL_d$) ($\bar{v}$ = 0.7674 $cm^3\ g^{-1}$). Left: SV continuous c(s) distribution at 60 krpm (s = 1.098 S, $s_{20,w}$ = 1.144 S, f/f0 = 1.232, Mw = 9898 Da, 3.0x monomer mass at a 95% confidence level). Right: SE data and fitted curves from 44 – 60 krpm for a single-species model (Mw = 10039 Da, 3.0x monomer mass, 95% confidence limits 9837.0 – 10172.1 Da). Conditions: 20 °C, PBS, pH 7.4. SV and SE experiments were conducted at 100 µM concentrations.

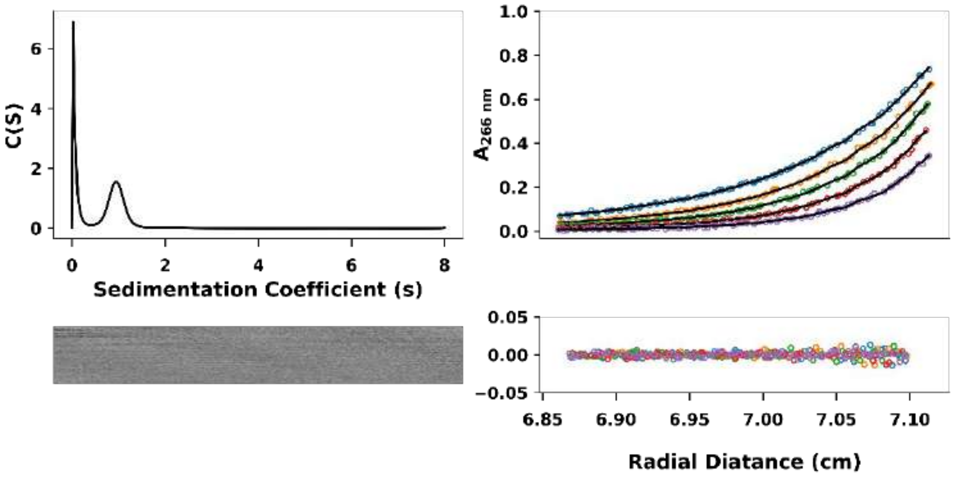


**Figure S4.2:** AUC data and fits (top) with residuals (bottom) for PK-2 ($\bar{v}$ = 0.7806 $cm^3\ g^{-1}$). Left: SV continuous c(s) distribution at 60 krpm (s = 0.949 S, $s_{20,w}$ = 0.991 S, f/f0 = 1.32, Mw = 9757 Da, 3.0x monomer mass at a 95% confidence level). Right: SE data and fitted curves from 44 – 60 krpm for a single-species model (Mw = 8932 Da, 2.6x monomer mass, 95% confidence limits 8725.6 – 9624.7 Da). Conditions: 20 °C, PBS, pH 7.4. SV and SE experiments were conducted at 100 µM concentrations.

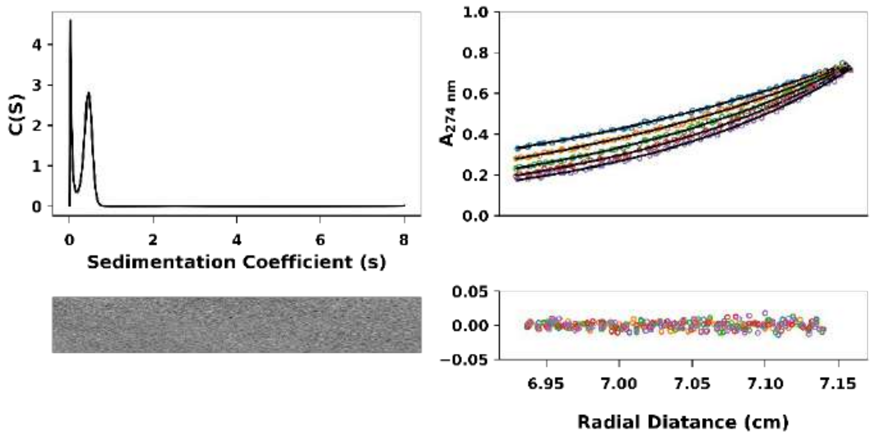


**Figure S4.3:** AUC data and fits (top) with residuals (bottom) for PK-3 ($\bar{v}$ = 0.7820 $cm^3\ g^{-1}$). Left: SV continuous c(s) distribution at 60 krpm (s = 0.444 S, $s_{20,w}$ = 0.463 S, f/f0 = 1.341, Mw = 3219 Da, 1.1x monomer mass at a 95% confidence level). Right: SE data and fitted curves from 44 – 60 krpm for a single-species model (Mw = 2972 Da, 1.0x monomer mass, 95% confidence limits 2833.6 – 3000.0 Da). Conditions: 20 °C, PBS, pH 7.4. SV and SE experiments were conducted at 100 µM concentrations.

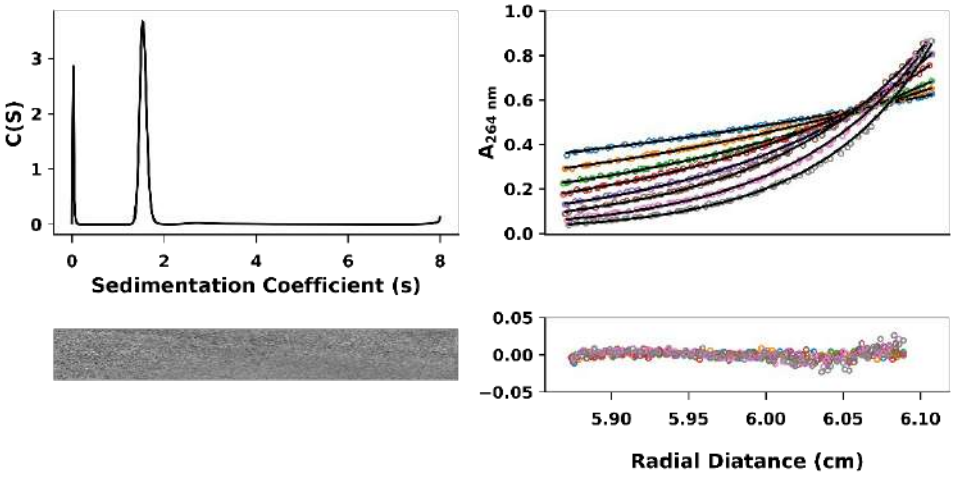


**Figure S4.4:** AUC data and fits (top) with residuals (bottom) for PK-4 ($3_{10}$HD) ($\bar{v}$ = 0.7967 $cm^3\ g^{-1}$). Left: SV continuous c(s) distribution at 60 krpm (s = 1.547 S, $s_{20,w}$ = 1.619 S, f/f0 = 1.23, Mw = 20760 Da, 6.2x monomer mass at a 95% confidence level). Right: SE data and fitted curves from 44 – 60 krpm for a single-species model (Mw = 21298 Da, 6.3x monomer mass, 95% confidence limits 21220.5 – 21534.2 Da). Conditions: 20 °C, PBS, pH 7.4. SV and SE experiments were conducted at 100 µM concentrations.

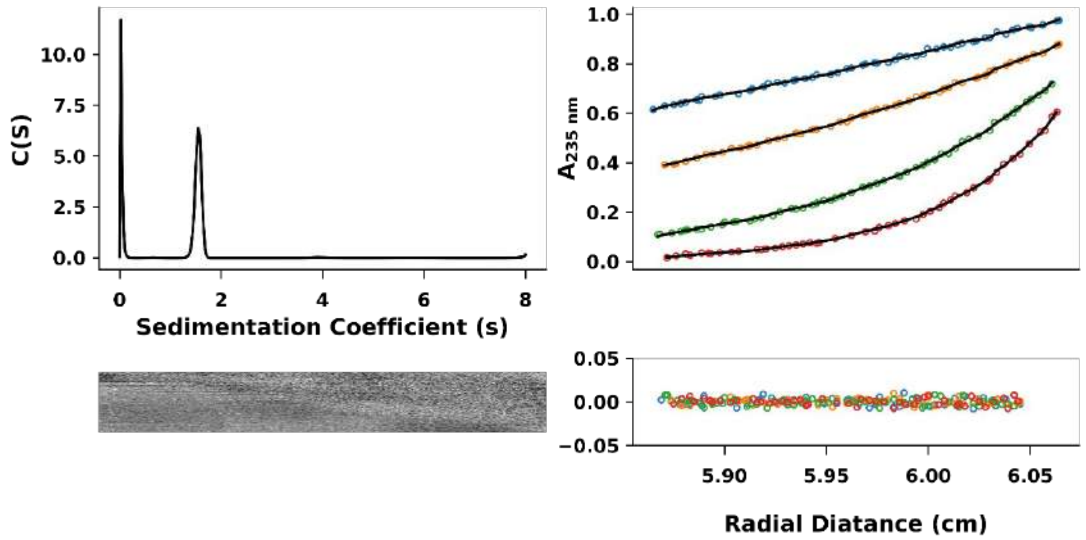


**Figure S4.5:** AUC data and fits (top) with residuals (bottom) for PK-5 (D-$3_{10}$HD) ($\bar{v}$ = 0.7957 $cm^3$ $g^{-1}$). Left: SV continuous c(s) distribution at 60 krpm (s = 1.552 S, $s_{20,w}$ = 1.624 S, f/f0 = 1.49, Mw = 27474 Da, 8.1x monomer mass at a 95% confidence level). Right: SE data and fitted curves from 15 – 40 krpm for a single-species model (Mw = 23334.9 Da, 6.9x monomer mass, 95% confidence limits 21818.5 – 24789.9 Da). Conditions: 20 °C, PBS, pH 7.4. SV and SE experiments were conducted at 50 µM concentrations. Shown here the data at speeds 15K, 20K, 30K and 40K rpm.

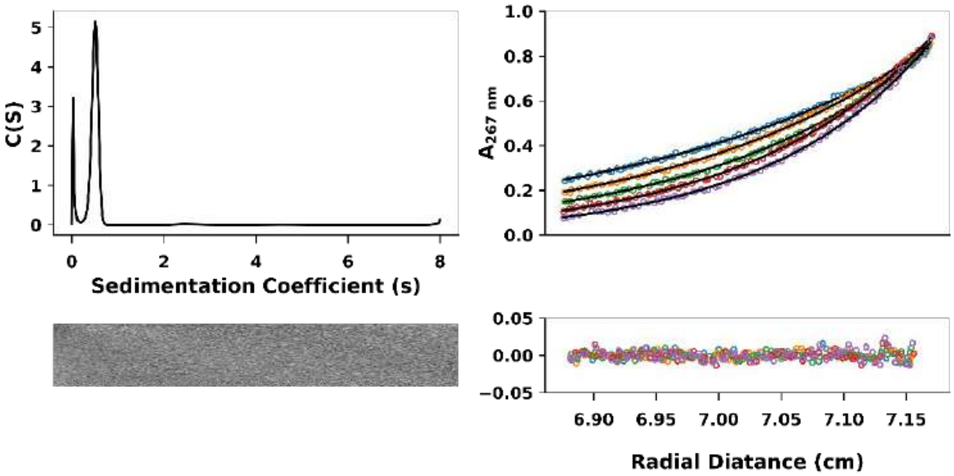


**Figure S4.6:** AUC data and fits (top) with residuals (bottom) for PK-6 ($\bar{v}$ = 0.7527 $cm^3$ $g^{-1}$). Left: SV continuous c(s) distribution at 60 krpm (s = 0.496 S, $s_{20,w}$ = 0.516 S, f/f0 = 1.26, Mw = 2800 Da, 0.9x monomer mass at a 95% confidence level). Right: SE data and fitted curves from 44 – 60 krpm for a single-species model (Mw = 2875 Da, 1.0x monomer mass, 95% confidence limits 2868.9 – 2944.4 Da). Conditions: 20 °C, PBS, pH 7.4. SV and SE experiments were conducted at 100 µM concentrations.

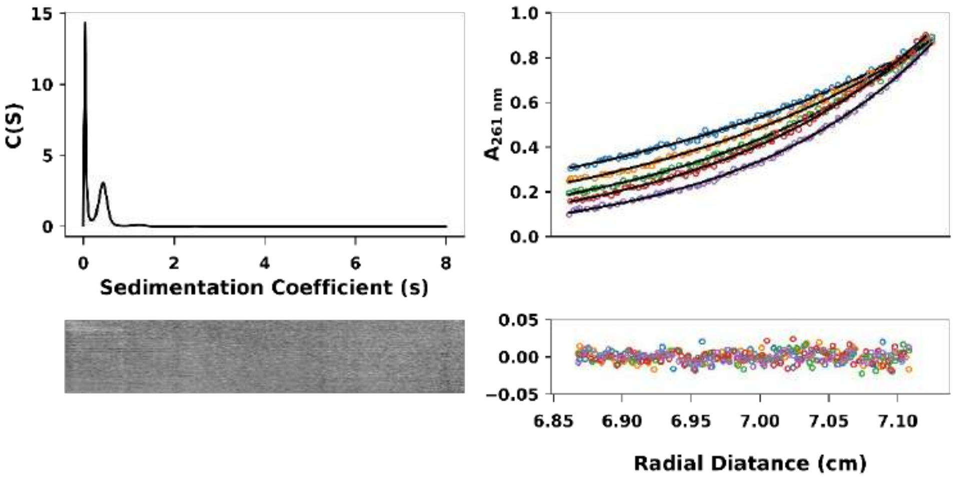


**Figure S4.7:** AUC data and fits (top) with residuals (bottom) for PK-7 ($\bar{v}$ = 0.7696 cm$^3$ g$^{-1}$). Left: SV continuous c(s) distribution at 60 krpm (s = 0.435 S, $s_{20,w}$ = 0.454 S, f/f0 = 1.58, Mw = 3650 Da, 1.2x monomer mass at a 95% confidence level). Right: SE data and fitted curves from 44 – 60 krpm for a single-species model (Mw = 2972 Da, 1.0x monomer mass, 95% confidence limits 2899.1 – 3030.6 Da). Conditions: 20 °C, PBS, pH 7.4. SV and SE experiments were conducted at 100 µM concentrations.

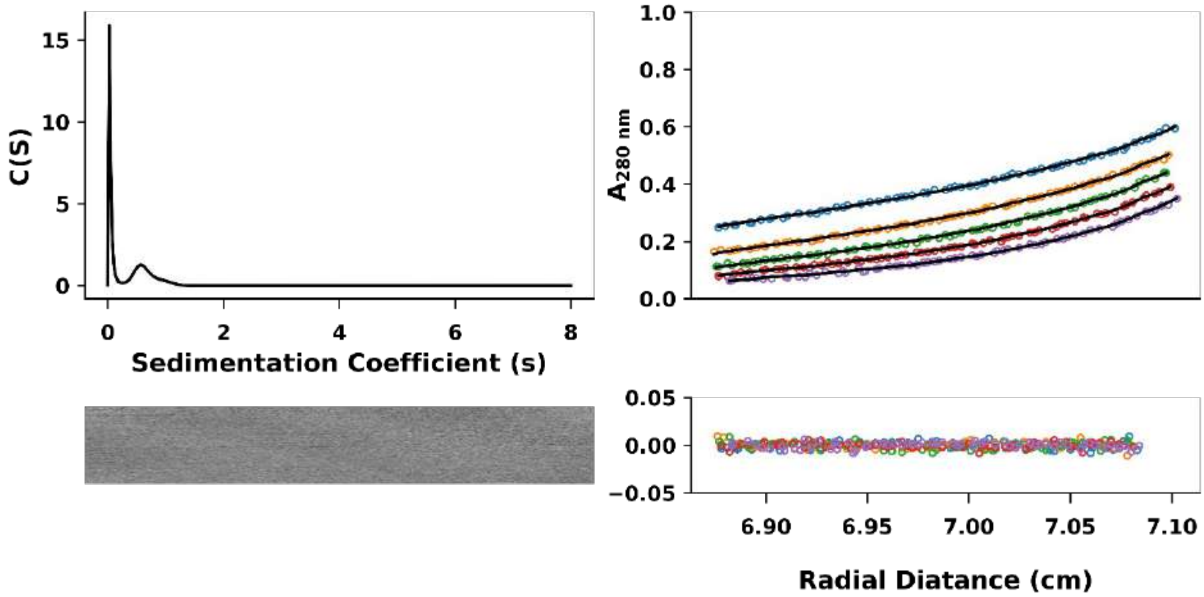


**Figure S4.8:** AUC data and fits (top) with residuals (bottom) for PK-8 ($\bar{v}$ = 0.7696 cm$^3$ g$^{-1}$). Left: SV continuous c(s) distribution at 60 krpm (s = 0.661 S, $s_{20,w}$ = 0.691 S, f/f0 = 1.349, Mw = 6628 Da, 2.0x monomer mass at a 95% confidence level). Right: SE data and fitted curves from 44 – 60 krpm for a single-species model (Mw = 3097.6 Da, 0.9x monomer mass, 95% confidence limits 2663.9 – 3966.5 Da). Conditions: 20 °C, PBS, pH 7.4. SV and SE experiments were conducted at 100 µM concentrations.

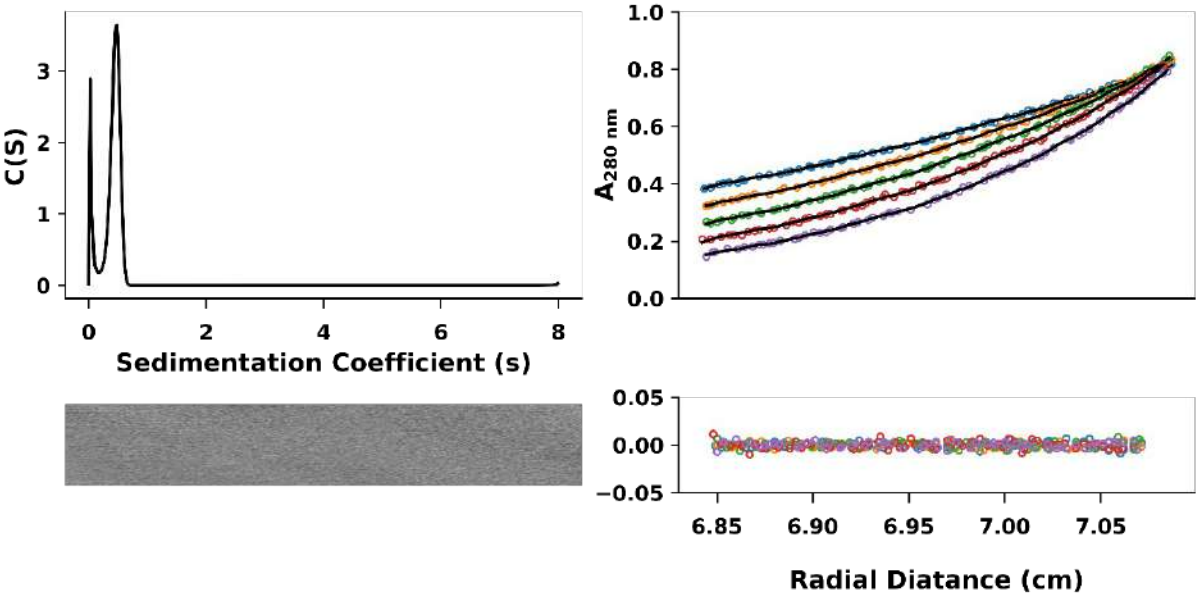


**Figure S4.9:** AUC data and fits (top) with residuals (bottom) for PK-9 ($\bar{\nu}$ = 0.7833 $cm^3$ $g^{-1}$). Left: SV continuous c(s) distribution at 60 krpm (s = 0.447 S, $s_{20,w}$ = 0.467 S, f/f0 = 1.298, Mw = 3130 Da, 1.0x monomer mass at a 95% confidence level). Right: SE data and fitted curves from 44 – 60 krpm for a single-species model (Mw = 3771.9 Da, 1.2x monomer mass, 95% confidence limits 3251.2 – 4379.3 Da). Conditions: 20 °C, PBS, pH 7.4. SV and SE experiments were conducted at 100 µM concentrations.

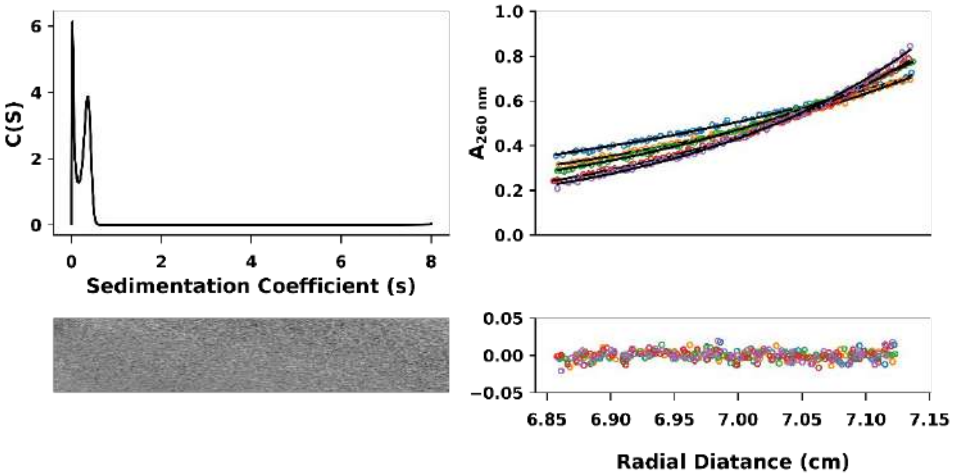


**Figure S4.10:** AUC data and fits (top) with residuals (bottom) for PK-10 ($\bar{\nu}$ = 0.7872 $cm^3$ $g^{-1}$). Left: SV continuous c(s) distribution at 60 krpm (s = 0.338 S, $s_{20,w}$ = 0.353 S, f/f0 = 1.23, Mw = 1955 Da, 1.0x monomer mass at a 95% confidence level). Right: SE data and fitted curves from 44 – 60 krpm for a single-species model (Mw = 2275.7 Da, 1.1x monomer mass, 95% confidence limits 2201.5 – 2331.8 Da). Conditions: 20 °C, PBS, pH 7.4. SV and SE experiments were conducted at 100 µM concentrations.

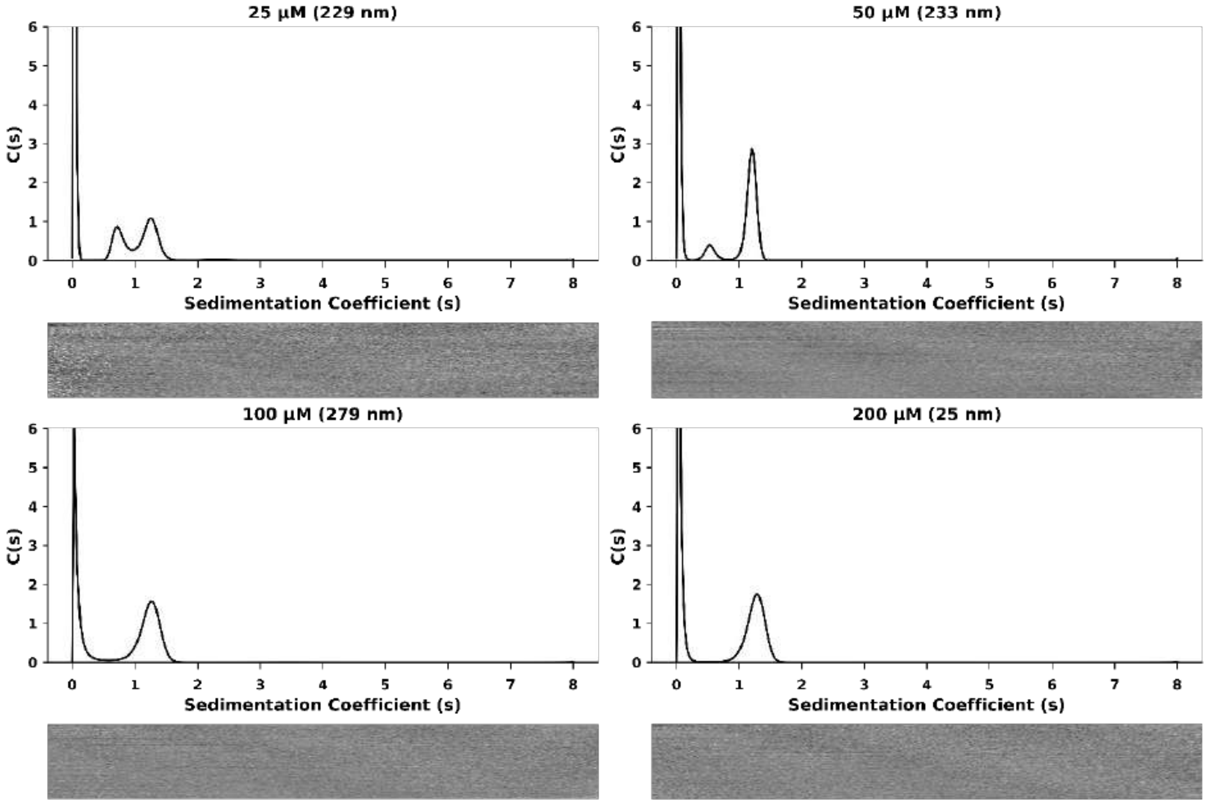


**Figure S4.11:** AUC sedimentation-velocity data for PK-12 at different concentrations. Data and fits (top) and residuals (bottom) ($\bar{v}$ = 0.7932 cm$^3$ g$^{-1}$). Each experiment was performed at 60 krpm. Top-left: Two peaks; s = (0.755, 1.241) S, $s_{20,w}$ = (0.789, 1.297) S, f/f0 = 1.346 and mw = (7840, 16517) Da (2.9, 6.1)x monomer mass at 95% confidence level. Top-right: Two peaks; s = (0.541, 1.199) S, $s_{20,w}$ = (0.566, 1.254) S, f/f0 = 1.182 and mw = (3920, 12922) Da (1.5, 4.8)x monomer mass at 95% confidence level. Bottom-Left: s = 1.225 S, $s_{20,w}$ = 1.281 S, f/f0 = 1.247 and mw = 14450 Da (5.3x monomer mass) at 95% confidence level. Bottom-right: Two peaks; s = 1.262 S, $s_{20,w}$ = 1.319 S, f/f0 = 1.254 and mw = 15231 Da (5.6x monomer mass) at 95% confidence level. Residuals (bottom of each panel) are shown as a bitmap in which the greyscale shade indicates the difference between the fit and raw data, where black and white colours represent data that have extreme deviations from the fit. PBS (pH 7.4) was used as buffer.

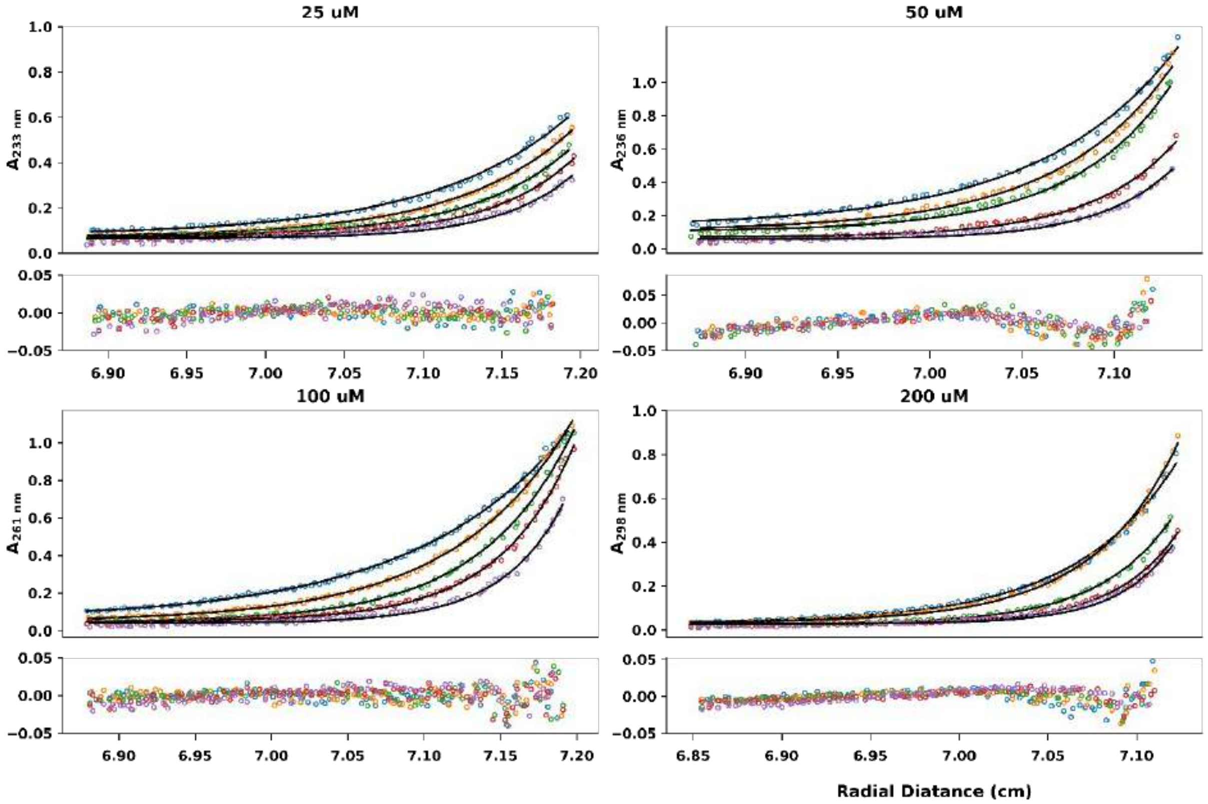


**Figure S4.12:** AUC sedimentation-equilibrium data for PK-12 at different concentrations. Data and fitted curves are from 44 – 60 krpm for a single-species model. Top-left: Concentration = 25 µM, Mw = 9669 Da, 3.6x monomer mass, 95% confidence limits 9439.8 – 9859.7 Da. Top-right: Concentration = 50 µM, Mw = 11405 Da, 4.2x monomer mass, 95% confidence limits 1111.3 – 11599.2 Da. Bottom-left: Concentration = 100 µM, Mw = 12852 Da, 4.8x monomer mass, 95% confidence limits 12676.6 – 12924.6 Da. Bottom-right: Concentration = 200 µM, Mw = 14928 Da, 5.6x monomer mass, 95% confidence limits 14731.7 – 15126.1 Da. Conditions: 20 °C, PBS, pH 7.4.

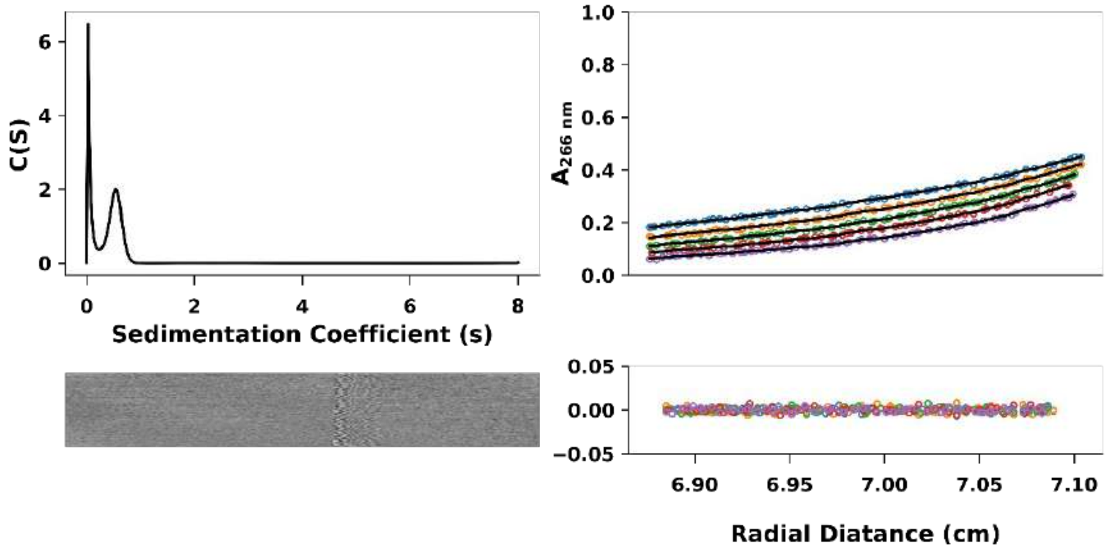


**Figure S4.13:** AUC data and fits (top) with residuals (bottom) for PK-13 ($3_{10}$HD-A) ($\bar{\nu}$ = 0.7720 $cm^3$ $g^{-1}$). Left: SV continuous c(s) distribution at 60 krpm (s = 0.550 S, $s_{20,w}$ = 0.573 S, f/f0 = 1.449, Mw = 4617 Da, 1.4x monomer mass at a 95% confidence level). Right: SE data and fitted curves from 44 – 60 krpm for a single-species model (Mw = 4371.6 Da, 1.3x monomer mass, 95% confidence limits 3016.9 – 4934.7 Da). Conditions: 20 °C, PBS, pH 7.4. SV and SE experiments were conducted at 100 µM concentrations.

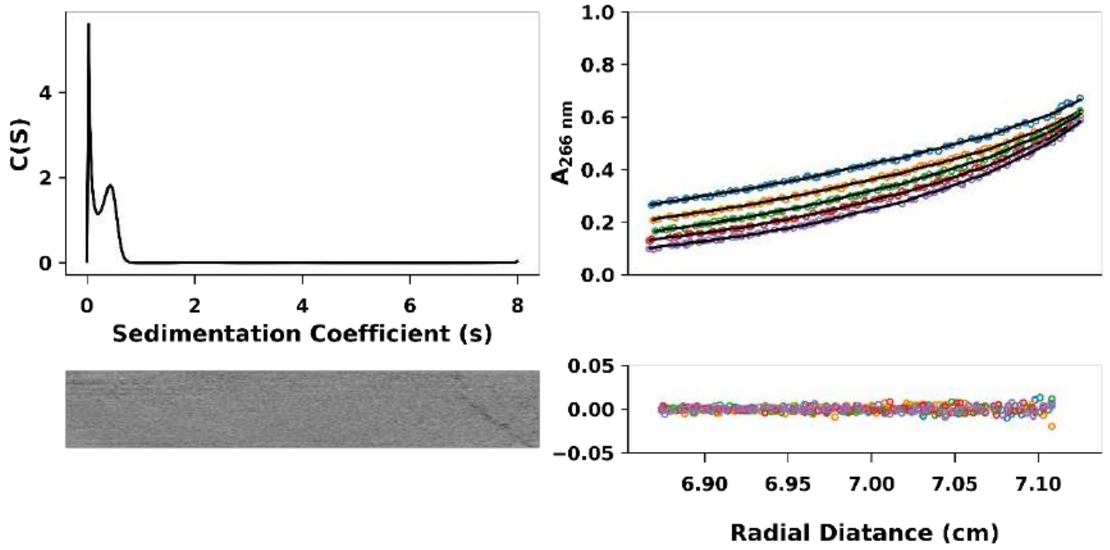


**Figure S4.14:** AUC data and fits (top) with residuals (bottom) for PK-14 ($3_{10}$HD-B) ($\bar{\nu}$ = 0.8820 $cm^3$ $g^{-1}$). Left: SV continuous c(s) distribution at 60 krpm (s = 0.418 S, $s_{20,w}$ = 0.440 S, f/f0 = 1.292, Mw = 3892 Da, 1.2x monomer mass at a 95% confidence level). Right: SE data and fitted curves from 44 – 60 krpm for a single-species model (Mw = 3044.8 Da, 0.9x monomer mass, 95% confidence limits 2585.1 – 3406.1 Da). Conditions: 20 °C, PBS, pH 7.4. SV and SE experiments were conducted at 100 µM concentrations.

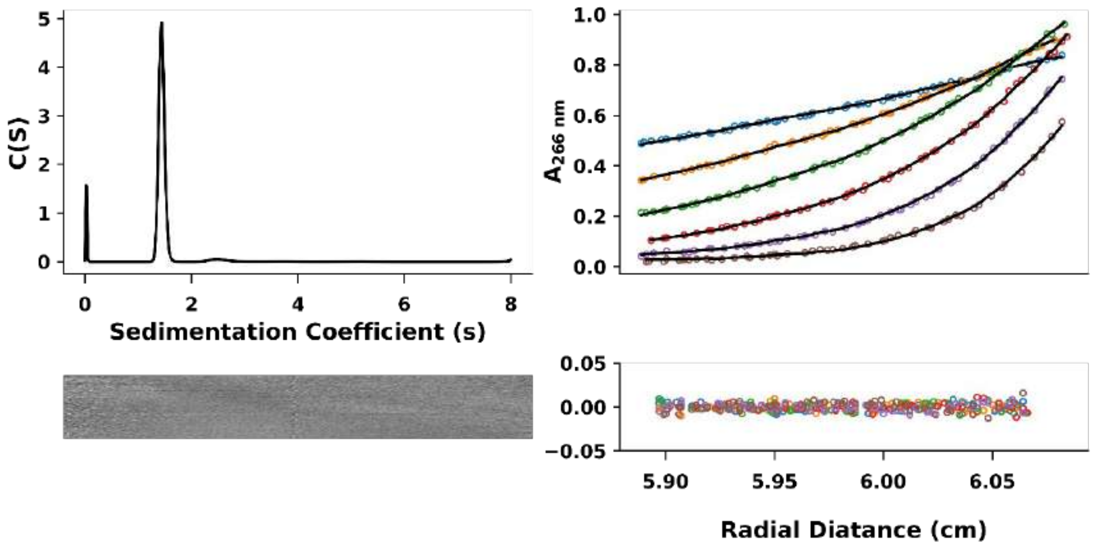


**Figure S4.15:** AUC data and fits (top) with residuals (bottom) for the heteromeric $3_{10}$HD-AB system (PK-13 ($3_{10}$HD-A) plus PK-14 ($3_{10}$HD-B)) (average $\bar{v}$ = 0.7967 $cm^3$ $g^{-1}$). Left: SV continuous c(s) distribution at 60 krpm (s = 1.441 S, $s_{20,w}$ = 1.508 S, f/f0 = 1.282, Mw = 19792 Da, 5.9x monomer mass at a 95% confidence level). Right: SE data and fitted curves from 18 – 48 krpm for a single-species model (Mw = 18030.8 Da, 5.4x monomer mass, 95% confidence limits 17567.6 – 18160.7 Da). Conditions: 20 °C, PBS, pH 7.4. SV and SE experiments were conducted at 100 µM concentrations.

**Section 5: DPH-binding Analyses**

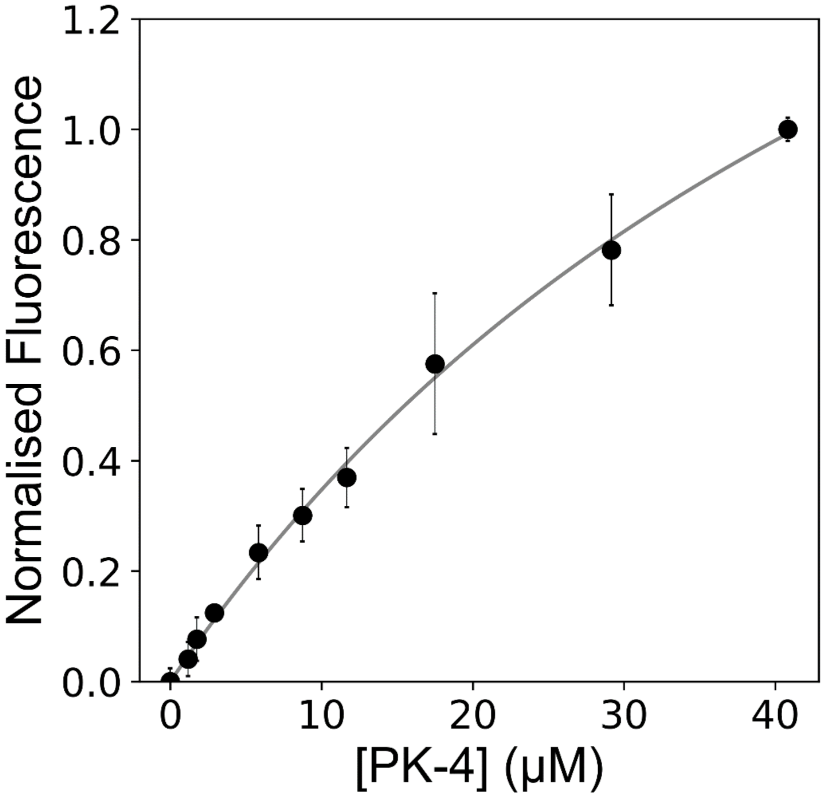


**Fig S5.1:** Titration curve for DPH with PK-4 ($3_{10}$HD). The data did not show any specific binding of DPH suggesting the $3_{10}$HD assembly has a consolidated hydrophobic core.

**Section 6: Structural Analyses of the $3_{10}$-helix bundle formed by PK-5 (D-$3_{10}$HD; PDB id 7qdi)**

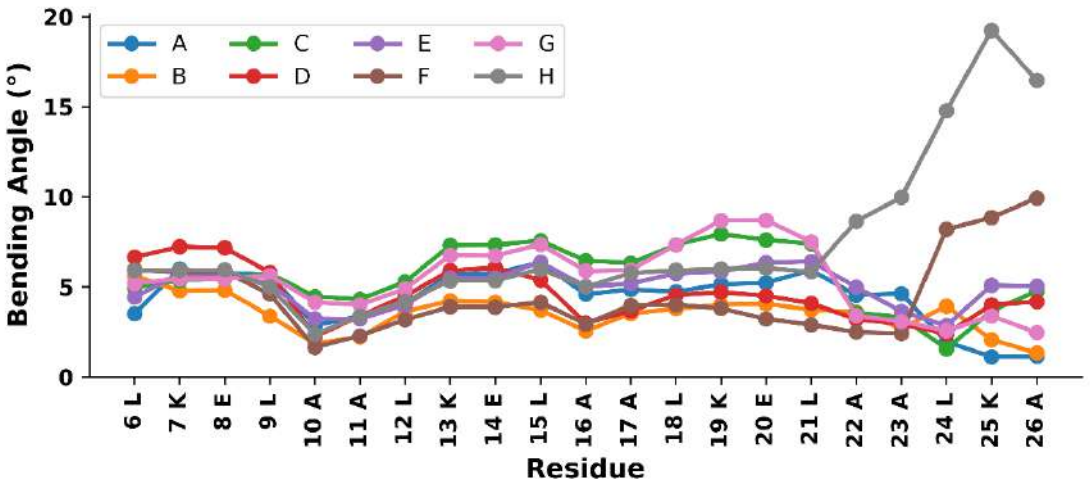


**Fig S6.1:** Bending angle between two consecutive turns of helix according to HELANAL-Plus. The bending angle is assigned to the residue that is common to the two turns. For example, the angle between two turns comprising residues 3-6 and 6-9 is assigned to residue 6 and so on. This analysis indicates that the helices of PK-5 (D-$3_{10}$HD) curve smoothly.

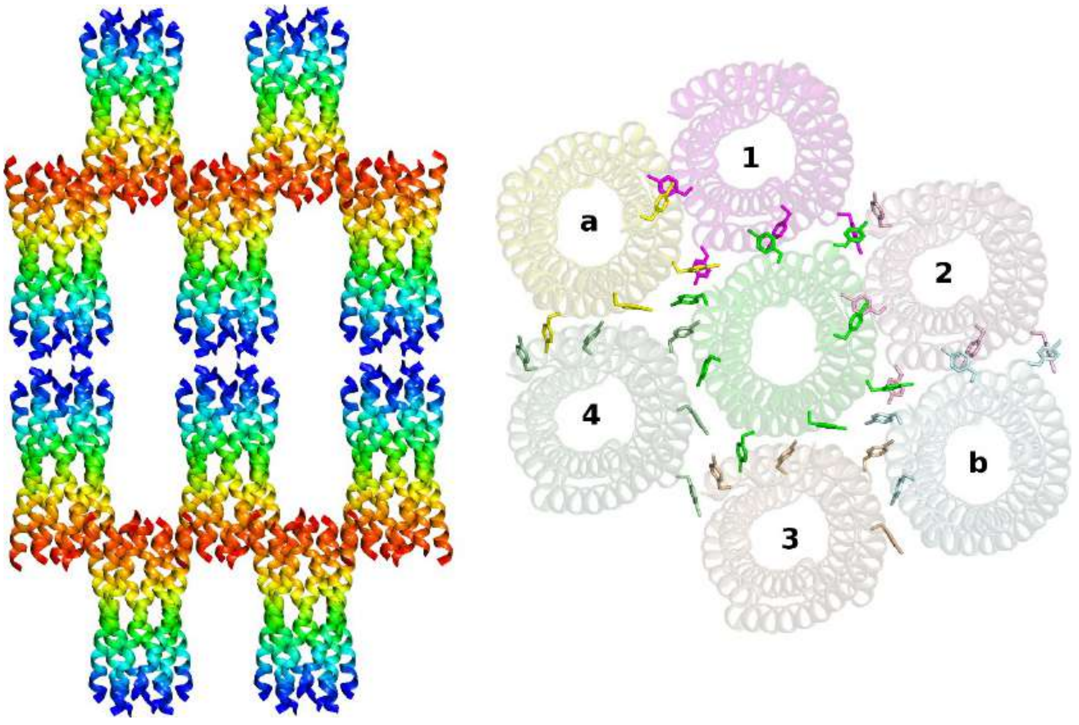


**Fig S6.2:** Packing of $3_{10}$-helix bundles in the unit cell of 7qdi. Left: The *N* termini are coloured blue and the *C*-termini red. Right: Groove made by helical bundles 1, 2, 3 and 4. Helical bundles a and b are at the same level to that of the middle assembly (coloured in green). D-Br-Phe residues from each helix of different assemblies are shown as sticks to show the aromatic-aromatic interactions.

**Section 7: Tables**

| Residues | $3_{10}$ helices | | | α helices | π helices | Total in Database |
|---|---|---|---|---|---|---|
| | < 6 Residues (1) | ≥ 6 Residues (2) | (1)+(2) | | | |
| **ALA** | 7072 | 710 | 7782 | 88184 | 744 | 188892 |
| **ARG** | 3848 | 465 | 4313 | 46622 | 588 | 117031 |
| **ASN** | 3944 | 354 | 4298 | 23448 | 561 | 98628 |
| **ASP** | 6656 | 601 | 7257 | 36741 | 632 | 136202 |
| **CYS** | 827 | 71 | 898 | 7692 | 143 | 28093 |
| **GLN** | 3425 | 347 | 3772 | 36260 | 397 | 85946 |
| **GLU** | 7409 | 763 | 8172 | 67235 | 966 | 151270 |
| **GLY** | 4777 | 506 | 5283 | 24081 | 582 | 163898 |
| **HIS** | 2159 | 256 | 2415 | 15090 | 343 | 54112 |
| **ILE** | 2340 | 328 | 2668 | 45167 | 931 | 129331 |
| **LEU** | 7063 | 1038 | 8101 | 92052 | 1422 | 213196 |
| **LYS** | 4815 | 603 | 5418 | 48409 | 635 | 127750 |
| **MET** | 1228 | 157 | 1385 | 16199 | 230 | 39352 |
| **PHE** | 3496 | 420 | 3916 | 30117 | 815 | 94467 |
| **PRO** | 5639 | 467 | 6106 | 14955 | 69 | 105276 |
| **SER** | 6277 | 593 | 6870 | 35026 | 439 | 137490 |
| **THR** | 3146 | 313 | 3459 | 31280 | 613 | 124535 |
| **TRP** | 1585 | 189 | 1774 | 11459 | 247 | 33857 |
| **TYR** | 3095 | 387 | 3482 | 25879 | 745 | 82638 |
| **VAL** | 2602 | 310 | 2912 | 46807 | 1094 | 158657 |
| **Total** | 81403 | 8878 | 90281 | 742703 | 12196 | 2270621 |

| **Parameters** | | | | | | |
|---|---|---|---|---|---|---|
| | **$3_{10}$ helices** | | **α helices** | | **π helices** | |
| | **Calculated** | **Ideal** | **Calculated** | **Ideal** | **Calculated** | **Ideal** |
| **Rise per Residue (Å)** | 1.88±0.26 | 2.00 | 1.50±0.13 | 1.50 | 1.31±0.21 | 1.10 |
| **Residues per Turn** | 3.28±0.25 | 3.00 | 3.65±0.13 | 3.60 | 4.17±0.21 | 4.30 |
| **Radius (Å)** | 2.02±0.19 | 1.90 | 2.31±0.10 | 2.30 | 2.61±0.16 | 2.80 |
| **ϕ (°)** | -66.33±33.09 | | -64.77±11.00 | | -80.13±24.28 | |
| **ψ (°)** | -15.11±27.82 | | -39.48±10.65 | | -41.78±17.39 | |

**Table S7.1:** Total amino acids counts and general information for various helices in protein structures from the PDB. Total number of such helices is given in parentheses. Mean and standard deviation for different helical parameters of helix-types along with the corresponding ideal values are also listed. Distribution of Ramachandran angles (ϕ, ψ) are shown in **Figure S1.2**.

| **Peptide** | **Crystallization Condition** | **Molecular dimensions screen** | **Beamline** |
|---|---|---|---|
| PK-1 (CCTri-TypeN-$L_aL_d$) | 0.09 M Halogens; 0.1 M Buffer System 2; 7.5 30 % v/v Precipitant Mix 2 | Morpheus® | I24 |
| PK-5 (D-$3_{10}$HD) | 0.1 MIB 6; 20% v/v Peg 500 MME | PACT PremierTM | I03 |
| PK-10 +PK-11 | 0.1 M MIB 8.0; 25 % w/v PEG 1500 | PACT PremierTM | I04-1 |

**Table S7.2:** List of crystallisation conditions, parent screen names and beamline used to collect dataset.

| | PK-1 (CCTri-TypeN-$L_aL_d$) | PK-5 (D-$3_{10}$HD) | PK-10+PK-11 |
|---|---|---|---|
| **PDB Code** | 7QDK | 7QDI | 7QDJ |
| **Data collection** | | | |
| Wavelength | 0.9190 | 0.9204 | 0.9119 |
| Resolution range | 19.25 - 1.41 (1.461 - 1.41) | 29.57 - 2.34 (2.42 - 2.34) | 15.69 - 1.44 (1.47 - 144) |
| Space group | P 31 | P 21 2 21 | C 1 2/c 1 |
| Cell dimensions | | | |
| a, b, c (Å) | 22.2 22.2 128.3 | 33.2 45.0 194.2 | 26.9 19.4 55.6 |
| α, β, γ (°) | 90.0 90.0 120.0 | 90.0 90.0 90.0 | 90.0 96.6 90.0 |
| Total reflections | 136068 (9449) | 139093 (13791) | 66985 (4227) |
| Unique reflections | 13655 (1357) | 12818 (1056) | 5148 (305) |
| Multiplicity | 10.0 (7.0) | 10.9 (11.3) | 13.0 (13.9) |
| Completeness (%) | 99.56 (98.76) | 97.13 (83.68) | 97.1 (94.8) |
| Mean I/sigma(I) | 21.57 (3.32) | 11.97 (1.98) | 41.0 (18.9) |
| Wilson B-factor | 16.87 | 42.54 | - |
| $R_{merge}$ | 0.06754 (0.5036) | 0.1138 (1.06) | 0.032 (0.098) |
| $R_{meas}$ | 0.07111 (0.5432) | 0.1196 (1.11) | 0.034 (0.101) |
| $R_{pim}$ | 0.0221 (0.1999) | 0.0358 (0.327) | 0.009 (0.027) |
| $CC_{1/2}$ | 0.999 (0.897) | 0.999 (0.824) | 1.0 (0.99) |
| $CC^*$ | 1 (0.973) | 1 (0.951) | - |
| **Refinement** | | | |
| Reflections used in refinement | 13614 (1356) | 12639 (1056) | 4677 (351) |
| Reflections used for R-free | 656 (80) | 593 (60) | 230 |
| $R_{work}$ | 0.1569 (0.1986) | 0.1890 (0.2809) | 0.1295 (0.1500) |
| $R_{free}$ | 0.1977 (0.2074) | 0.2390 (0.3342) | 0.1679 (0.1940) |
| $CC_{work}$ | 0.965 (0.864) | 0.948 (0.666) | - |
| $CC_{free}$ | 0.956 (0.923) | 0.935 (0.696) | - |
| Number of non-hydrogen atoms | 811 | 2070 | 202 |
| macromolecules | 691 | 552 | 160 |
| ligands | 29 | 1450 | 15 |
| solvent | 91 | 68 | 27 |
| Protein residues | 90 | 94 | 19 |
| RMS bond lengths (Å) | 0.013 | 0.018 | 0.011 |
| RMS bond angles (°) | 1.64 | 3.21 | 1.407 |
| Ramachandran favoured (%) | 100.00 | - | 100.00 |
| Ramachandran allowed (%) | 0.00 | - | 0.00 |
| Ramachandran outliers (%) | 0.00 | - | 0.00 |
| Rotamer outliers (%) | 1.89 | 0.00 | 0.00 |
| Clash score | 4.79 | 4.45 | 0.00 |
| Average B-factor | 21.82 | 51.78 | 10.00 |
| macromolecules | 19.78 | 44.33 | 9.44 |
| ligands | 31.85 | 54.11 | 9.28 |
| solvent | 34.06 | 62.55 | 18.98 |
| Number of TLS groups | 3 | 8 | 1 |

**Table S7.3:** Merging and refinement statistics for all X-ray crystal structures presented in this paper. Highest-resolution shell parameters are shown in parentheses. $R_{free}$ represents the R-factor calculated from 5% of reflections that were not used during refinement.
